# Temperature-dependent herbivore nutritional traits affect population dynamics and persistence

**DOI:** 10.1101/2025.10.28.685165

**Authors:** David M. Anderson, Francis Fruin Laid-Low, Mary I. O’Connor

## Abstract

Herbivores can vary in their relative requirements for nutrients and carbon, with important consequences for consumer-resource dynamics and nutrient cycling. The nutrient (phosphorus) and carbon requirements of herbivore growth have both been found to increase at temperature extremes, yet the consequences of these thermal responses for herbivore-autotroph population dynamics have not been explored. Here, we develop and analyze a stoichiometrically-explicit, temperature-dependent model of herbivore-autotroph systems to evaluate how these two thermal responses of herbivore nutritional traits affect population responses to temperature and phosphorus supply. We find contrasting effects of increased phosphorus or carbon demands at temperature extremes. Increases in herbivore phosphorus demands relatively strongly restricts the range of temperatures at which the herbivore population persists and slightly decreases the stability of population dynamics at high phosphorus supplies. In contrast, increases in the carbon requirements of herbivore growth slightly decreases the temperature range supporting herbivore persistence and stabilizes the system under phosphorus enrichment. These contrasting effects of herbivore nutrient and carbon demands reflect differences in the ecological contexts at which they influence herbivore performance. Increased nutrient requirements impair herbivore productivity near the herbivore persistence boundary where autotrophs are abundant and nutrient-poor, whereas high carbon demands decrease herbivore productivity near the stability boundary where autotrophs are at low density and nutrient-rich. The thermal response of herbivore nutritional traits may therefore be an important factor influencing population responses to warming and nutrient enrichment.

## Introduction

Temperature and nutrient supply strongly influence physiology and population dynamics. Temperature constrains organismal metabolism and energetics (Robinson et al. 1983, Raven and Geider 1988, Gillooly et al. 2001), and, in doing so, affects ecological processes including growth, mortality, and consumption (Eppley 1971, Brown et al. 2004, Dell et al. 2011). Effects of nutrient availability on ecological processes are mediated by organismal stoichiometry, because biomass production and maintenance require a balance of chemical elements (Sterner and Elser 2002, Prater et al. 2024). Despite substantial work on the individual-level effects of temperature or nutrients on organismal metabolism and stoichiometry, integrating and scaling these effects up to population and community dynamics remains a major challenge (Allen and Gillooly 2009, Cross et al. 2015, Meunier et al. 2024, O’Connor et al. 2025, Tumolo et al. 2025).

Efforts to integrate individual energetics and stoichiometry have focused on the nutritional requirements of consumer growth (Doi et al. 2010, Cross et al. 2015). Consumers require energy stored in carbon-carbon bonds and nutrients in the form of chemical elements (e.g., phosphorus, P) to produce new biomass. Mismatches between consumer requirements and resource stoichiometry can inhibit resource assimilation, energy acquisition and ultimately reduce consumer growth rate (Urabe and Watanabe 1992, Frost et al. 2006, Prater et al. 2024). The resource stoichiometry (C:X ratio, where “X” is an element such as P) that distinguishes between carbon- (or energy-) and nutrient-limited growth is often referred to as the Threshold Elemental Ratio (hereafter, *TER_C:X_* or simply “nutritional requirements,” Urabe and Watanabe 1992, Frost et al. 2006). The *TER_C:X_* is an important trait of individual consumers that affects demographic rates and controls the flow of energy and nutrients through consumer biomass.

Temperature can affect consumer nutritional traits by altering the uptake and assimilation of carbon and nutrients. Recent studies have identified two patterns of responses of nutritional (phosphorus) requirements (*TER_C:P_*) to temperature. First, Ruiz et al. (2020) found that the *TER_C:P_* exhibits a U-shaped relationship to temperature in the cladoceran *Daphnia magna*. This increase in carbon demands at thermal extremes arose because carbon acquisition (ingestion) rates were strongly decreased relative to carbon requirements for respiration (Schmitz and Rosenblatt 2017, Ruiz et al. 2020). Second, Laspoumaderes et al. (2022) found that the *TER_C:P_* is a unimodal function of temperature in four aquatic herbivore species (including *D. magna,* the protist *Oxyrrhis marina*, copepod *Acartia tonsa*, and crab *Carcinus maenus*). Increases in phosphorus demands at thermal extremes occurred because ingested phosphorus was converted into biomass less efficiently than carbon; this difference in growth efficiencies also produced increases in the C:P ratio of herbivore biomass at temperature extremes (Laspoumaderes et al. 2022). These two patterns could be parts of a more complex N-shaped thermal response of the *TER_C:P_*, although this relationship has only been observed in one species (*C. maenus*, Laspoumaderes et al. 2022). Monotonic increases or decreases in the *TER_C:P_* with warming have also been found (e.g., Persson et al. 2011, Wojewodzic et al. 2011, Malzahn et al. 2016), but these studies are thought to have sampled an insufficient temperature range to capture the U- (or hump-)shaped response (Schmitz and Rosenblatt 2017, Ruiz et al. 2020). Despite these advances in understanding of the thermal response of herbivore nutrient traits, we know little about their implications for population dynamics.

Herbivore nutritional traits have important consequences for herbivore-autotroph population dynamics (Elser and Urabe 1999, Urabe et al. 2002). Herbivores and autotrophs differ in stoichiometric flexibility: autotroph stoichiometry is plastic and sensitive to environmental conditions (Droop 1974), whereas herbivore stoichiometry is homeostatically regulated (Andersen and Hessen 1991, Sterner and Elser 2002). These differences affect population dynamics (Andersen et al. 2004). For example, herbivore populations can collapse when autotroph populations are abundant but low in nutritional quality (Sommer 1992, Sterner et al. 1998, Diehl et al. 2022, Frost et al. 2025). Herbivore population growth can be facilitated by increased autotroph nutritional quality that arises from external changes (e.g., nutrient inputs, Frost et al. 2025; but see Nessel et al. 2021) and/or intrinsic feedbacks with herbivore populations (e.g., via nutrient excretion, Sterner 1990, Urabe et al. 2002, or a reduction in autotroph density that concentrates nutrients in autotroph biomass, Sommer 1992, Urabe 1995, Hall et al. 2007a). Herbivore nutritional traits affect these dynamics by determining both herbivore sensitivity to poor food quality, and the impact of herbivores on autotroph nutritional contents (Andersen 1997, Elser and Urabe 1999).

The thermal traits of herbivore populations can also influence the dynamics of herbivore-autotroph systems. Analyses of bioenergetic consumer-resource (herbivore-autotroph) models have shown that temperatures that support high herbivore growth and survival rates generally yield large energy fluxes from autotroph to herbivore and correspondingly unstable population dynamics, and temperature changes that strongly reduce herbivore productivity (e.g., to temperatures above or below that which is optimal for growth) can reduce energy fluxes, increase stability, and lead to herbivore extinction (Rall et al. 2010, Uszko et al. 2017, Vinton and Vasseur 2022). Studies that incorporate nutrients and stoichiometric constraints into temperature-dependent consumer-resource models have shown that decreased nutrient supply can decrease the thermal ranges supporting stable population dynamics and herbivore persistence (Uszko et al. 2017), and that flexible autotroph stoichiometry can further decrease the thermal breadth of herbivore persistence (Sentis et al. 2022). However, it is unclear how temperature-dependent shifts in nutritional traits (Ruiz et al. 2020, Laspoumaderes et al. 2022) may influence these patterns of population responses to temperature and nutrients.

Here, we evaluate the effects of temperature-dependent herbivore nutritional traits on the response of autotroph-herbivore systems to temperature and nutrient supply. We develop a stoichiometrically-explicit, temperature-dependent model describing the coupled dynamics of autotroph and herbivore populations. Our model builds on work by Sentis et al. (2022), which integrates general theory describing stoichiometric constraints on consumer-resource interactions (Andersen 1997, Loladze et al. 2000, Muller et al. 2001, Andersen et al. 2004, Hall et al. 2007b, Hall 2009) with the model of Uszko et al. (2017) that considers interactive effects of temperature and resources on trophic interactions. Specifically, we modify this modeling framework to allow the herbivore’s *TER_C:P_* to exhibit unimodal or U-shaped responses to temperature following empirical findings by Laspoumaderes et al. (2022) and Ruiz et al. (2020), respectively. Our objective is to evaluate how the thermal response of herbivore nutritional traits impacts the sensitivity of three important community properties – 1) herbivore persistence, 2) system stability and population fluctuations, and 3) the trophic interaction strength (Gilbert et al. 2014) – to temperature and nutrient supply. We find that thermal variation in herbivore nutritional traits alters dynamic feedbacks between autotroph stoichiometry and herbivore population density, and that these changes can impact the structure and dynamics of the community.

## Model and Methods

### Model Development

We develop a model describing how temperature and phosphorus supply influence interactions between an herbivore population with a fixed body stoichiometry and an autotroph population with a variable stoichiometry. Body stoichiometry is expressed as the Phosphorus:Carbon (P:C) ratio (the inverse of the C:P ratio, which had been considered in the Introduction). To test the effects of temperature-dependent herbivore nutritional traits on community properties – herbivore persistence, system stability, and interaction strength – we contrast the following three scenarios: 1) herbivores have constant nutritional requirements and body stoichiometry across temperatures, 2) herbivores require more nutritionally dense (i.e., high P:C ratio) autotrophs to maximize growth rate and are themselves less nutritionally dense (i.e., have a lower P:C ratio) at thermal extremes (Laspoumaderes et al. 2022), and 3) herbivores require more carbon-rich (i.e., low P:C ratio) autotrophs to maximize growth rate at thermal extremes and have constant body stoichiometry (Ruiz et al. 2020).

The model consists of two differential equations describing the coupled dynamics of the autotroph (*A*, gC m^−3^) and herbivore biomass density (*Z*, gC m^−3^), with an algebraic equation that determines the autotroph Phosphorus:Carbon stoichiometry (*Q_A_*, gP gC^−1^; also referred to here simply as phosphorus or nutrient contents) (Fig. 1, Table S1):

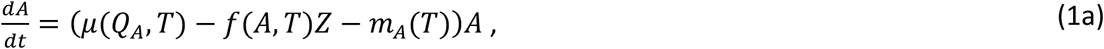

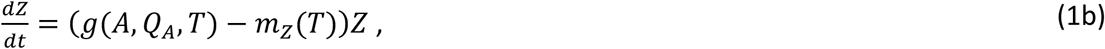

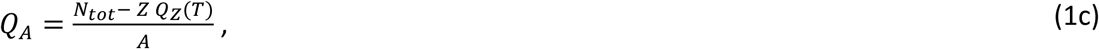

where *μ*(*Q_A_*, *T*) describes how the autotroph’s specific rate of carbon production (or growth rate, gC gC^−1^d^−1^) depends on its’ phosphorus contents and temperature *T* (°C), *f*(*A*, *T*) describes how the herbivore’s grazing rate (m^3^ d^−1^gC^−1^) depends on autotroph population density and temperature, *g*(*A*, *Q_A_*, *T*) captures the effects of autotroph population density, stoichiometry, and temperature on herbivore specific growth rate (gC gC^−1^ d^−1^), and *m_A_*(*T*) and *m*_Z_(*T*) are temperature-dependent rates of biomass loss to mortality for autotrophs and herbivores, respectively (gC gC^−1^d^−1^). This model tracks units of both carbon and phosphorus in biomass, in order to consider the effects of stoichiometry (i.e., phosphorus:carbon ratios, *Q_A_* and *Q*_Z_(*T*)) and metabolism on consumer-resource dynamics.

**Fig. 1.**
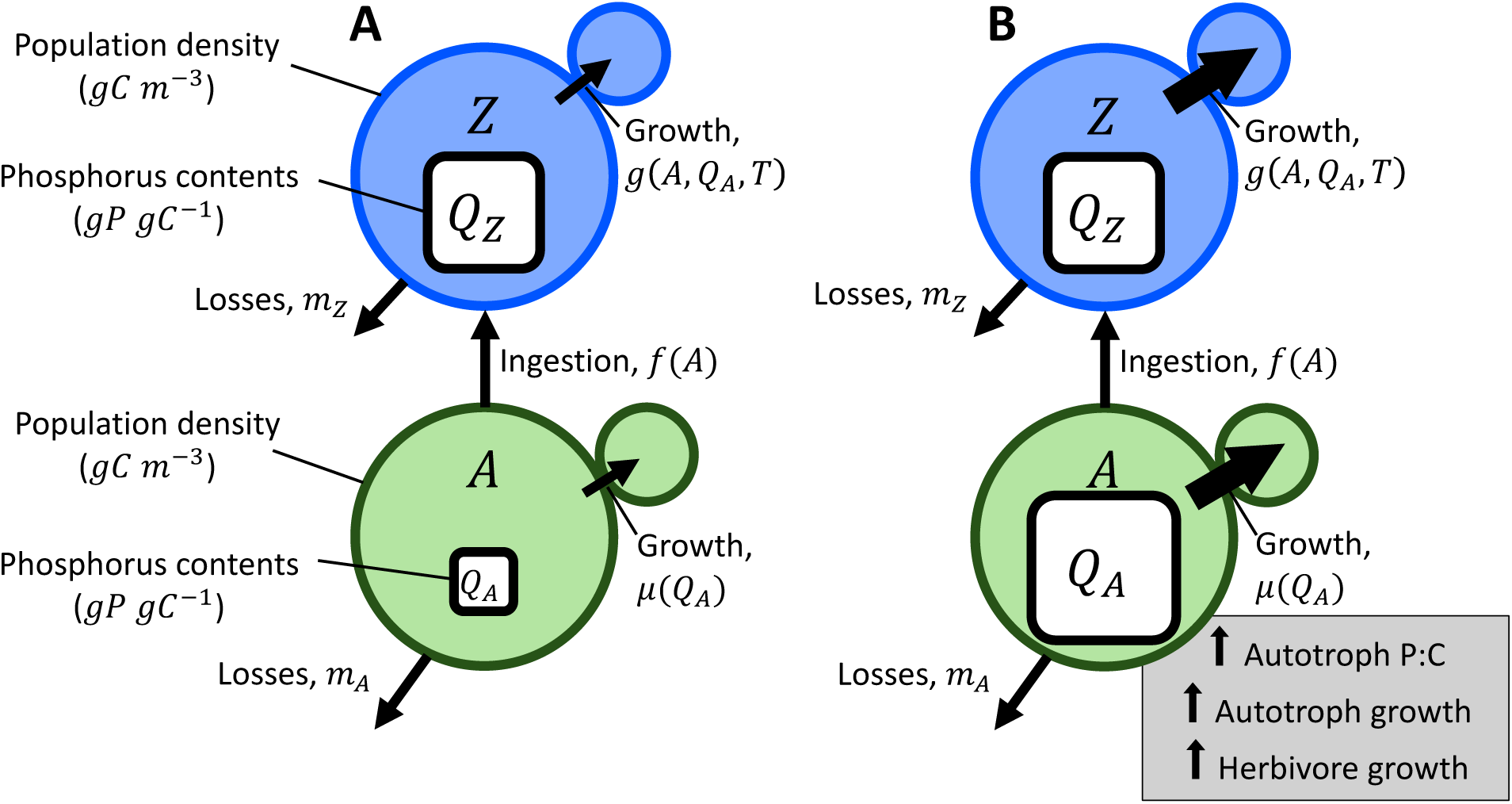
Conceptual diagram of the stocks and flows of biomass in the model. **A** The green and blue circles represent biomass densities of the autotroph *A* and herbivore *Z*, respectively, the inset white squares depict phosphorus:carbon stoichiometry, and the arrows show the flows of biomass in the system (note that growth, or the production of new biomass, is represented here by the flow of biomass from the larger circle to the smaller circle). Panel **B** shows that increases in the phosphorus contents of the autotroph, which may arise from increases in phosphorus availability *N_tot_* or decreases in herbivore or autotroph population density (see Eqn. 1c), leads to an increase in the growth rate of both the autotroph population (Eqn. 4a) and herbivore population (via improved phosphorus growth rate, Eqn. 2c).

The autotroph phosphorus content (Eqn. 1c, *Q_A_*) is variable, and depends on how the total phosphorus availability, *N_tot_* (gP m^−3^), is divided into herbivore and autotroph biomass. This model follows from two key assumptions regarding phosphorus cycling: 1) the system is closed for phosphorus, and 2) fluxes of phosphorus from other pools within the system (i.e., P embedded in dead biomass and in inorganic form) back into autotroph biomass happens instantaneously (Andersen 1997). These two assumptions generally do not qualitatively alter model outcomes; Diehl 2007 and Diehl et al. 2022 contrasted this model structure to one explicitly resolving phosphorus cycling through detritus and inorganic pools and found no qualitative difference. Under these assumptions, all available phosphorus is incorporated into either autotroph or herbivore biomass. The phosphorus contents of autotrophs *Q_A_* can therefore be calculated by rearranging an equation describing mass-balance constraints on phosphorus (Eqn. 1c, Andersen 1997, Loladze et al. 2000, Andersen et al. 2004): the total phosphorus in the system *N_tot_* (gP m^−3^) is the sum of all phosphorus incorporated into herbivore biomass (i.e., *Z Q*_Z_(*T*), where *Q*_Z_(*T*) is the temperature-dependent herbivore stoichiometry in gP gC^−1^) and autotroph biomass (i.e., *A Q_A_*).

The herbivore specific growth rate is limited either by the specific rate of carbon growth *g_C_*(*A*, *T*) (gC gC^−1^d^−1^) or the specific rate of phosphorus growth *g_P_*(*A*, *Q_A_*, *T*) (gP gP^−1^d^−1^) (Hall et al. 2007b):

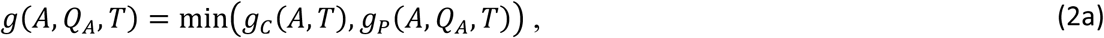

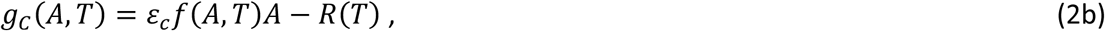

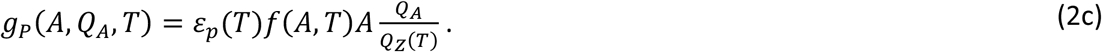

The carbon growth rate *g_C_*(*A*, *T*) is determined by the product of carbon ingestion rate (*f*(*A*, *T*)*A*, gC gC^−1^d^−1^) and the carbon conversion efficiency *ε_c_* (gC gC^−1^), minus the carbon lost to respiration *R*(*T*) (gC gC^−1^d^−1^). Similarly, the rate of phosphorus production *g* (*A*, *Q* , *T*) is determined by the phosphorus ingestion rate 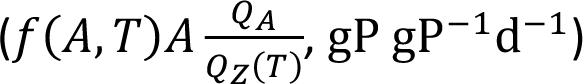 and the temperature-dependent phosphorus conversion efficiency *ε_p_*(*T*) (gP gP^−1^). Importantly, the threshold elemental ratio (*TER_P:C_*) of herbivore growth occurs at the autotroph food stoichiometry *Q_A_* that yields co-limitation between carbon and phosphorus growth (i.e., when *g_C_*(*A*, *T*) = *g_P_*(*A*, *Q_A_*, *T*)). The temperature dependence of the *TER_P:C_* will depend on the relative thermal sensitivities of ingestion, respiration, phosphorus conversion efficiency, and herbivore stoichiometry. We next specify these temperature dependencies, and we then describe how they combine to determine herbivore nutrient demands across temperatures.

The rates at which herbivores ingest autotrophs and respire carbon vary with temperature *T* (°C) and/or autotroph population density (expressed as carbon density) as:

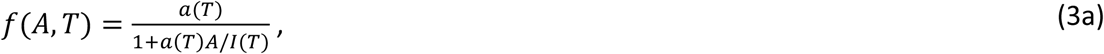

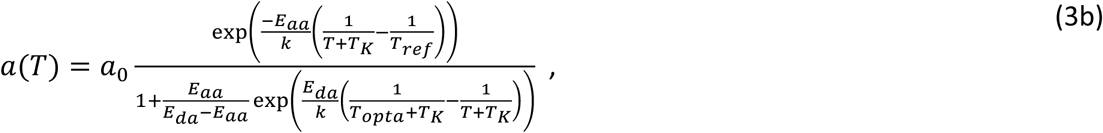

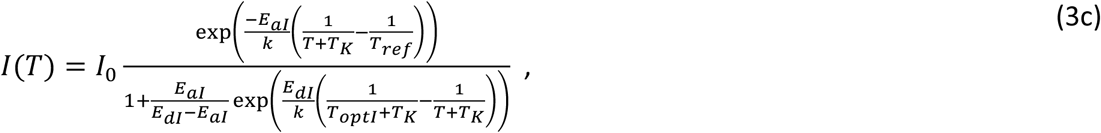

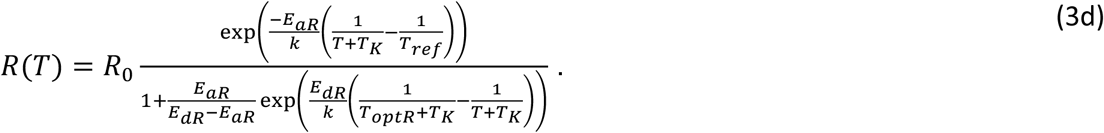

The clearance rate *a*(*T*) (m^3^ d^−1^gC^−1^), maximum ingestion rate *I*(*T*) (i.e., the inverse of the handling time, d^−1^), and respiration rate *R*(*T*) vary unimodally with temperature as described by Sharpe-Schoolfield functions (Schoolfield et al. 1981), with a maximum value at temperature *T_opta_*, *T_optI_*, or *T_optR_* and thermal sensitivity controlled by *E_aa_*, *E_aI_*, or *E_aR_* below the thermal optimum and *E_da_*, *E_dI_*, or *E_dR_* above the thermal optimum (note that Eqns. 3b-d rearrange the Sharpe-Schoolfield function to introduce a thermal optimum parameter). The parameter *k* = 8.62 ∗ 10^−5^ is the Boltzmann’s constant (eV K^−1^), *T_K_* = 273.15 is included to convert temperature *T* in °C to Kelvin, and *T_ref_* = 298.15 K is a reference temperature. These left-skewed, unimodal thermal responses are consistent with data for the temperature-dependence of the functional response (Englund et al. 2011, Uszko et al. 2017, Uiterwaal and DeLong 2020), including from Laspoumaderes et al. (2022) (Fig. S1A). The herbivore stoichiometry *Q*_Z_(*T*) and phosphorus conversion efficiency *ε_p_*(*T*) may exhibit a unimodal temperature-dependence following:

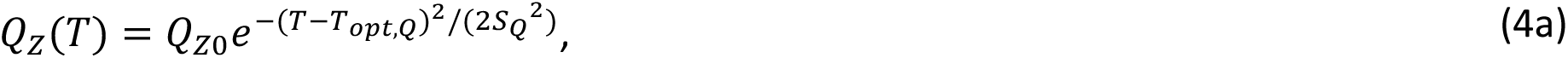

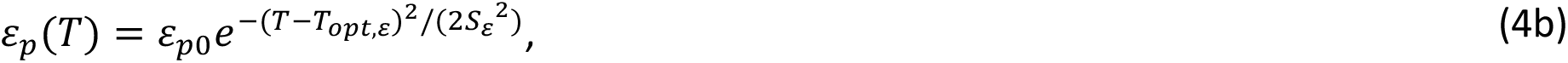

where *T_opt_*_,s_ and *T_opt_*_,Q_ give the temperatures maximizing these functions and *S*_s_ and *S*_Q_ determine the thermal breadths (Fig. 2C, Fig. S1). These symmetric functions describe data for the temperature-dependence of herbivore nutrient contents and demands from Laspoumaderes et al. (2022) reasonably well (see subsequent text and Fig. S1).

**Fig. 2.**
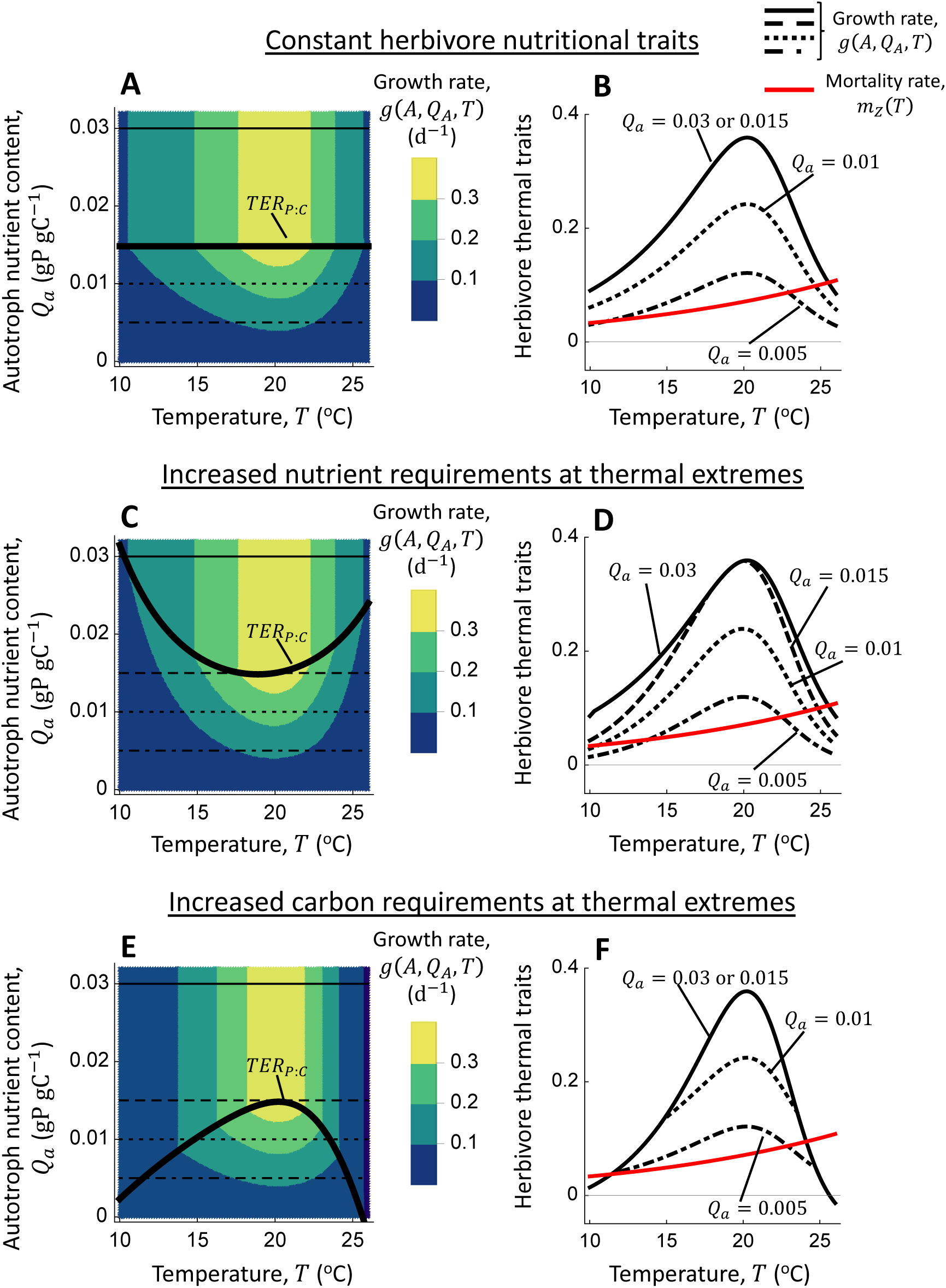
Effects of the temperature-dependence of herbivore nutrient (phosphorus) requirements (*TER_P_*_:*C*_(*T*), Eqn. 5) on growth rate (*g*(*A*, *Q_A_*, *T*), Eqns. 2). **A-B** Herbivore growth rate under the assumption that phosphorus requirements (*TER_P_*_:*C*_(*T*), solid black line in **A**) are insensitive to temperature. **C-D** Herbivore growth rate when phosphorus requirements (*TER_P_*_:*C*_(*T*), solid black line in **C**) increase towards warm and cold temperatures. **E-F** Herbivore growth rate when the *TER_P_*_:*C*_(*T*) (solid black line in **C**) is a hump-shaped function of temperature, such that carbon requirements are higher at temperature extremes. Panels **A**, **C**, and **E** show growth rate as a function of temperature and autotroph nutritional quality *Q_A_* when autotroph density is not limiting (*A* → ∞). The horizontal lines indicate the values of autotroph nutritional quality *Q_A_* considered in panels **B**, **D**, and **F**. In **B** and **F**, thermal performance curves for growth rate at *Q_A_* = 0.03 and *Q_A_* = 0.015 lie on top of one another. Note that our models track P:C ratios (*TER_P_*_:*C*_) whereas Laspoumaderes et al. (2022) and Ruiz et al. (2020) report C:P ratios (*TER_C_*_:*P*_) and, as such, the thermal responses here are inverted.

The herbivore’s nutritional requirements, *TER_P_*_:*C*_(*T*), vary with temperature following (Fig. 2):

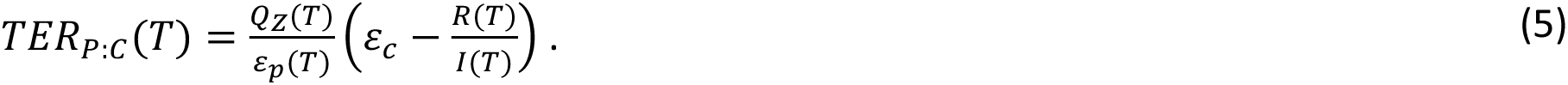

Eqn. (5) was derived by setting *g_C_*(*A*, *T*) = *g_P_*(*A*, *Q_A_*, *T*) while letting *A* → ∞ (as the *TER_P_*_:*C*_ is commonly assayed when food availability is not limiting, Frost et al. 2006, Ruiz et al. 2020, Laspoumaderes et al. 2022), and then solving for autotroph stoichiometry *Q_A_*. The three scenarios regarding the form of *TER_P_*_:*C*_(*T*) – including cases in which the *TER_P_*_:*C*_(*T*) is invariant, U-shaped, and hump-shaped across the temperature gradient – reflect differences in the temperature-dependencies of the underlying parameters and are illustrated in Fig. 2. The first scenario, in which nutrient demands *TER_P_*_:*C*_(*T*) are constant, arises when the herbivore stoichiometry *Q*_Z_ and phosphorus conversion efficiency *ε_p_* are independent of temperature (modeled here by setting *S*_s_ and *S*_Q_ to large numbers) and herbivore respiration and ingestion rates exhibit identical temperature-dependencies (i.e., *E_aI_* = *E_aR_*, *E_dI_* = *E_dR_*, and *T_optI_* = *T_optR_*, Fig. 2A-B, Fig. S1A-B).

Second, the U-shaped temperature dependence of *TER_P_*_:*C*_(*T*) observed by Laspoumaderes et al. (2022) is modeled here by allowing the herbivore body stoichiometry *Q*_Z_(*T*) and phosphorus conversion efficiency *ε_p_*(*T*) to exhibit unimodal temperature dependencies (Eqn. 4), while respiration and ingestion rates change in parallel across the temperature gradient (as in scenario 1). Note that the decrease in P:C ratios, *Q*_Z_(*T*), at temperature extremes leads to decreases in nutrient demands and hence a hump-shaped temperature-dependence of *TER_P_*_:*C*_(*T*). The U-shaped temperature-dependence of *TER_P_*_:*C*_(*T*) arises here due to a relatively strong reduction in phosphorus conversion efficiency *ε_p_*(*T*) at warm and cool temperatures (Fig. S1D; a U-shaped temperature response of carbon conversion efficiency *ε_c_* can also create a U-shaped *TER_P_*_:*C*_(*T*), but this inflates growth rate at low temperatures or produces a bimodal temperature response of herbivore growth rate, see Fig. S2). The herbivore growth rate is more sensitive to reductions in food nutritional quality at temperature extremes in this scenario (Fig. 2C-D).

Third, the hump-shaped temperature dependence of *TER_P_*_:*C*_(*T*) occurs when ingestion rates decline more strongly than respiration rates as temperatures diverge from the thermal optimum (i.e., *E_aI_* > *E_aR_* and *E_dI_* > *E_dR_*, Ruiz et al. 2020). As in the first scenario and consistent with data in Ruiz et al. (2020), herbivore stoichiometry *Q*_Z_ and phosphorus conversion efficiency *ε_p_* are temperature-independent. In this scenario, the herbivore growth rate tends to be carbon-limited at temperature extremes because carbon requirements for respiration are large relative to carbon acquisition (ingestion) rates (Fig. 2E-F).

The autotroph population’s specific growth rate increases as its phosphorus content *Q_A_* increases above its’ minimum requirements *Q_min_* (gP gC^−1^), up to a maximum temperature-dependent birth rate *μ*_∞_(*T*) (gC gC^−1^d^−1^, Droop 1974):

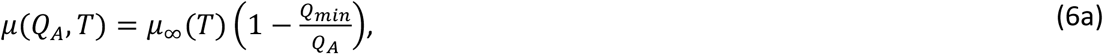

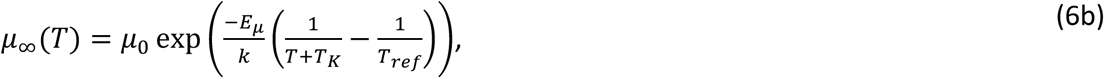

where the maximum birth rate *μ*_∞_(*T*) increases with temperature following the Boltzmann-Arrhenius function (Gillooly et al. 2001). The parameter *μ*_0_ is a scaling coefficient (d^−1^), *E_μ_* determines the sensitivity of birth rate to temperature (the apparent activation energy of metabolic reactions sustaining growth, eV^−1^), and *k*, *T_K_*, and *T_ref_* are constants defined above. Eqn. 6a-b assumes that temperature impacts the maximum growth rate *μ*_∞_, but not the minimum nutrient requirements *Q_min_*, consistent with past theoretical work (Sentis et al. 2022, Sauterey and Ward 2022, Bieg and Vasseur 2024) and empirical findings by Anderson et al. (2025).

Lastly, the mortality rates for the autotroph and herbivore, *m_A_*(*T*) and *m*_Z_(*T*) respectively, scale with temperature following the Boltzmann-Arrhenius function (Gillooly et al. 2001):

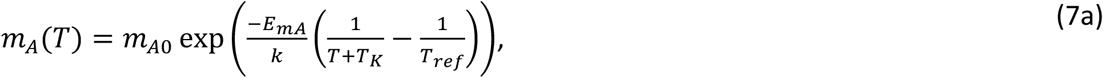

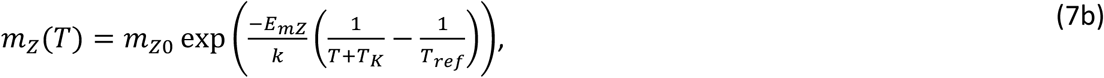

where *m_A_*_0_ and *m*_Z0_ are the scaling coefficients (d^−1^), and *E_mA_* and *E_m_*_Z_ determine the thermal sensitivities of these rates. This exponential temperature-dependence of mortality losses follows from the Metabolic Theory of Ecology (Brown et al. 2004) and is consistent with evidence from comparative analyses (Gillooly et al. 2001, McCoy and Gillooly 2008, Munch and Salinas 2009), and it is therefore commonly employed in temperature-dependent population models (Uszko et al. 2017, Sentis et al. 2022, Vinton and Vasseur 2022).

### Parameterization

The model (Eqn. 1-7) is parameterized based on data for the marine herbivorous zooplankton *Acartia tonsa* consuming the unicellular autotroph *Rhodomonas salina* (phytoplankton). For the scenarios “Constant herbivore nutritional traits” and “Increased nutrient requirements at thermal extremes,” datasets for *A. tonsa* maximum ingestion rate and respiration rate from Laspoumaderes et al. (2022) were pooled to estimate a common set of parameters controlling thermal sensitivities of these rates (i.e., assuming *E_aI_* = *E_aR_*, *E_dI_* = *E_dR_*, and *T_optI_* = *T_optR_* in Eqn. 3c-d; model fits are shown in Fig. S1A-B). For the scenario “Increased carbon requirements at thermal extremes” (Fig. 2), the temperature dependences of respiration rate were set at values lower than those of ingestion rate (*E_aR_* = 0.65 and *E_dR_* = 3) and the coefficient *R*_0_ was adjusted such that respiration rates at *T_optR_* were equivalent across scenarios (Fig. S1B). The herbivore clearance rate was estimated from a functional response experiment conducted on *A. tonsa* and *Rhodomonas spp.* at 18°C by Thor and Wendt (2010) (Fig. S1E), and we assumed that the thermal sensitivity of the clearance rate was identical to that of the maximum ingestion rate. Note that *A. tonsa* was found by Thor and Wendt (2010) to feed on *Rhodomonas* with a type 3 functional response. We study the type 2 functional response here because it is prevalent in natural systems (DeLong et al. 2025) and widely used in models of temperature-dependent consumer-resource dynamics (e.g., Synodinos et al. 2021, Sentis et al. 2022).

For the scenario “Increased nutrient requirements at thermal extremes,” parameters controlling the temperature dependence of herbivore stoichiometry were estimated by fitting Eqn. 4a to data in Laspoumaderes et al. (2022) (Fig. S1C). The carbon and phosphorus conversion efficiencies are *ε_c_* = 0.6 and *ε_p_*_0_ = 0.8, which are commonly observed values for herbivores (Frost et al. 2006, Halvorson et al. 2019), including *A. tonsa* (Thor and Wendt 2010). The thermal breadth of phosphorus conversion efficiency (*S*_s_) was estimated by fitting Eqn. 5 for *TER_P_*_:*C*_(*T*) to data from Laspoumaderes et al. (2022) (with other parameters fixed, Fig. S1D). The thermal optimum of phosphorus conversion efficiency is assumed to be equivalent to that for herbivore stoichiometry (i.e., *T_opt_*_,s_ = *T_opt_*_,Q_). The herbivore mortality rate was taken as 0.0485 at 15°C (based on estimates of non-predatory mortality of copepodites from Elliott and Tang 2011), with a thermal sensitivity of *E_m_*_Z_ = 0.54 as estimated by Hirst and Kiørboe (2002) for broadcast spawners like *A. tonsa*.

The temperature dependences of autotroph birth rate (Eqn. 6b) and mortality rate (Eqn. 7a) were estimated based on data for the net per-capita population growth rate of *R. salina* across temperatures from Ferreira et al. (2022) (Fig. S1F). The autotroph *Q_min_* was based on data for phosphorus limited *R. salina* from Meunier et al. (2012). Model fits were obtained using maximum likelihood methods, implemented with the Mathematica function “NonLinearModelFit[]”. Parameter values are provided in Table S1.

### Analysis

We manipulate the presence and form of thermal variation in herbivore nutrient requirements *TER_P_*_:*C*_(*T*) (Fig. 2), and evaluate how this affects community responses to temperature and phosphorus availability (*N_tot_*). The scenario “Constant herbivore nutrient requirements” (Fig. 2A-B) serves as a control for evaluating how increases in herbivore nutrient or carbon demands towards thermal extremes (Fig. 2C-D and Fig. 2E-F, respectively) influences outcomes. The scenarios “Increased nutrient requirements at thermal extremes” and “Increased carbon requirements at thermal extremes” are equivalent to the control case with constant nutrient requirements at *T_opt_*_,Q_ ≈ 18.9°C and *T_optI_* ≈ 20.2°C, respectively, and, as such, the differences between the scenarios are relevant as temperatures diverge from 18.9°C or 20.2°C (see also Fig. S1D). The “Constant herbivore nutrient requirements” scenario models low nutrient requirements (i.e., a low *TER_P_*_:*C*_) relative to the “Increased nutrient requirements at thermal extremes” scenario and low carbon requirements (i.e., a high *TER_P_*_:*C*_) relative to the “Increased carbon requirements at thermal extremes” scenario. In the Supplementary Materials, we also consider alternative *TER_P_*_:*C*_ values in the control scenario, including increased *TER_P_*_:*C*_ (due to impaired phosphorus conversion efficiency or enhanced carbon conversion efficiency) and decreased *TER_P_*_:*C*_ (due to improved phosphorus conversion efficiency or decreased carbon conversion efficiency).

We briefly summarize the community properties considered here and provide additional details in the Supplemental Materials section “Calculating community properties.” We identify three properties of the community and compare how they change over the temperature and nutrient supply gradients: 1) the herbivore persistence boundary, 2) the stability boundary, and 3) the strength of the trophic interaction. We solve for the sets of temperature and phosphorus availability that 1) separate herbivore persistence from extinction (i.e., herbivore persistence boundary; Eqn. S4, Fig. S3A-B) and 2) separate stable and unstable non-trivial equilibria (i.e., stability boundary; Fig. S3A). Because unstable equilibria lead to limit cycles with very low autotroph population densities that renders autotrophs (and the herbivores who depend on autotrophs) susceptible to extinction (Rosenzweig 1971), we also find temperatures and phosphorus supplies producing minimum autotroph population densities of 10^-2^ and 10^-4^.

Lastly, we evaluate the trophic interaction strength, defined here as the ratio of autotroph density in the absence of herbivores to the autotroph density in the presence of the herbivore (Eqn. S5) (Gilbert et al. 2014), across temperature and phosphorus supply gradients. This measure of interaction strength captures the flux of carbon from autotrophs to herbivore biomass and is therefore negatively related to system stability (Gilbert et al. 2014, Nilsson and McCann 2016), although it is not always monotonically related to stability metrics (e.g., the magnitude of the dominant eigenvalue, Uszko et al. 2017). High trophic interaction strengths occur when herbivores strongly reduce autotroph density relative to the density that autotrophs achieve in the absence of herbivory.

Equilibria, persistence boundaries, and stability boundaries were calculated using Mathematica (Version 12.1.1.0) and the EcoEvo package. We solved for equilibria analytically, and filtered out non-trivial equilibria that did not satisfy conservation of phosphorus (see Eqn. S3). We used root finding approaches to assess nutrient supplies required for persistence, destabilization of the equilibrium, and limit cycles producing minimum autotroph population densities of 10^-2^ and 10^-4^.

## Results

We observed qualitatively similar effects of temperature and nutrient supply on herbivore persistence, system stability, and interaction strength across the three scenarios (Fig. 3). The phosphorus supplies needed to support herbivore persistence or to destabilize population dynamics exhibited U-shaped responses to temperature, whereas the trophic interaction strength – the herbivore’s capacity to control autotroph population density – increased with increasing phosphorus supply and decreased as temperatures approached the herbivore persistence boundaries (Fig. 3A-C).

**Fig. 3.**
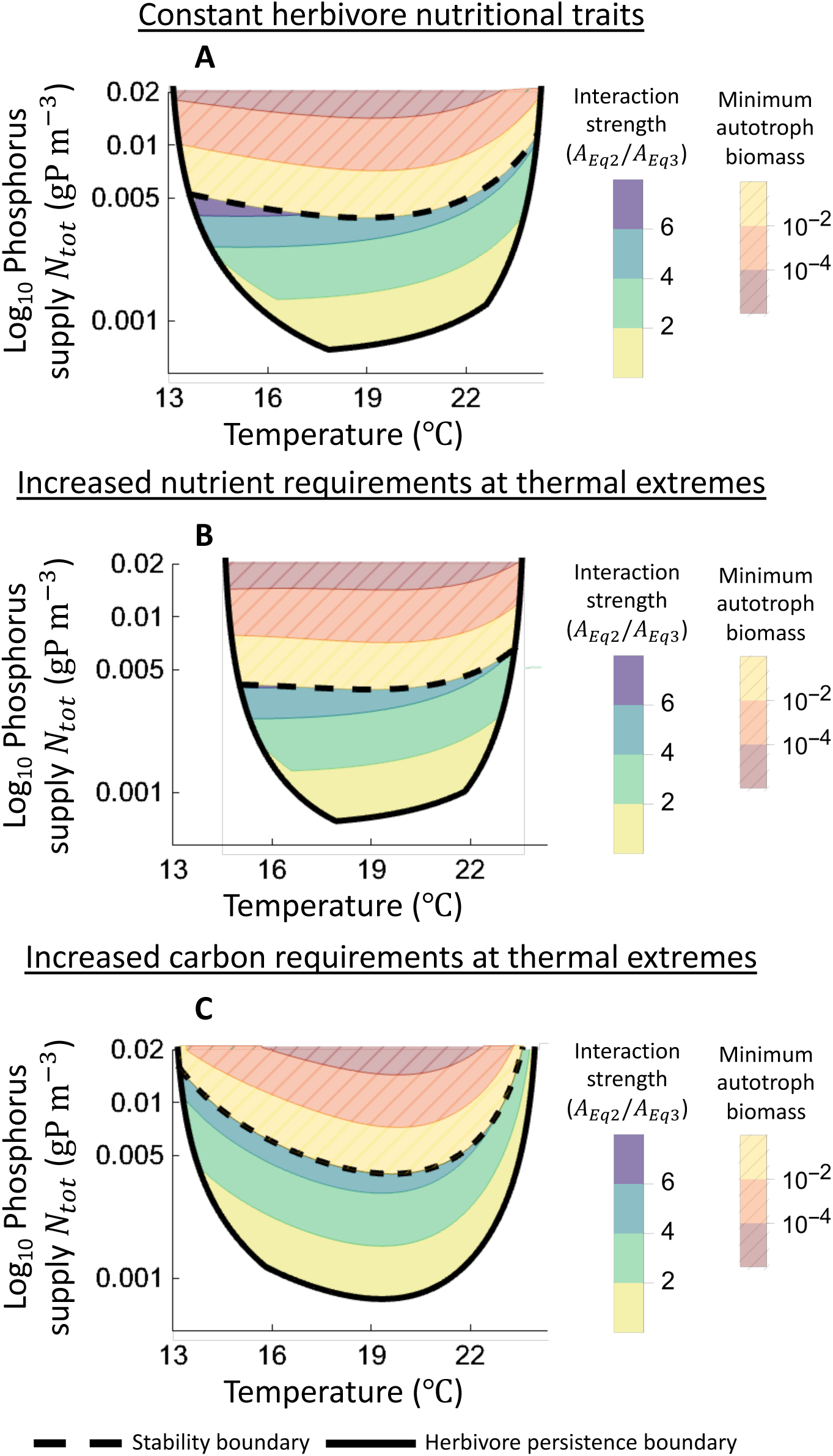
Effects of temperature and log_10_ transformed phosphorus supply on community properties when herbivores have constant nutrient requirements (A), increased nutrient demands at thermal extremes (B), and increased carbon requirements at thermal extremes (C). In each panel, herbivore persistence is supported above the solid black line, and the non-trivial equilibrium is unstable above the dashed black line. The trophic interaction strength – the equilibrium autotroph biomass in the presence of herbivores (denoted *A_Eq_*_2_) divided by the equilibrium autotroph biomass in the presence of herbivores (*A_Eq_*_3_; Eqn. S5) – is indicated by the shaded contours, and are only shown at phosphorus concentrations and temperatures at which herbivores persist with autotrophs at a stable equilibrium. The temperatures and phosphorus supplies producing limit cycles with minimum autotroph biomass densities less than or greater than 10^-2^ or 10^-4^ are indicated with the hatched shaded contours. See Fig. S4 for figures in which community properties for the three scenarios are displayed together to facilitate comparisons.

However, temperature-dependent shifts in herbivore nutritional traits produced quantitative changes in these patterns of community responses to temperature and nutrients. We focus on two sets of results, based on comparisons of each scenario with the control case in which herbivore nutrient requirements are constant (see also Fig. S4 for additional comparisons of community properties across scenarios). First, when thermal extremes cause herbivores to require more nutritionally dense autotrophs and have less nutritionally dense biomass, we find that, relative to the control scenario in which temperature does not affect nutritional traits: 1) the temperature range permitting herbivore persistence is restricted at all phosphorus concentrations *N_tot_* (contrast persistence boundaries in Fig. 3A and 3B; Fig. S4A), 2) the phosphorus supply required to destabilize the system decreases slightly (compare stability boundaries in Fig. 3A and 3B; Fig. S4B), and 3) the trophic interaction strength is reduced, but only near the persistence boundaries (contrast Fig. 3A and 3B; Fig. S4C).

Second, relative to the scenario in which herbivores have constant nutrient requirements, increases in carbon requirements at thermal extremes leads to 1) a small decrease in the thermal breadth of the herbivore persistence boundary (contrast persistence boundaries in Fig. 3A and 3C; Fig. S4A), 2) increases in the phosphorus supply at which the equilibrium is destabilized (especially at temperatures greater than 22°C and less than 16°C; compare stability boundaries in Fig. 3A and 3C; Fig. S4B), and 3) decreases in trophic interaction strengths (compare contours in Fig. 3A and 3C; Fig. S4C).

Additionally, we found that the consequences of increasing the herbivore’s relative requirements for phosphorus or carbon depended on the underlying mechanism (Supplementary Materials section “Effects of the herbivore *TER_P:C_* on community properties”). Increases in the herbivore’s phosphorus requirements that reflect an impaired phosphorus conversion efficiency increased the temperature range supporting herbivore persistence but did not influence stability (Fig. S5D), whereas increased herbivore P-requirements due to improved carbon conversion efficiency had little influence on herbivore persistence and destabilized population dynamics (Fig. S5C). Increases in the herbivore’s carbon requirements that emerge from improved phosphorus conversion efficiency increased the thermal breadth of the herbivore persistence boundary but did not impact stability (Fig. S5D), whereas increased herbivore carbon demands that reflect decreased carbon conversion efficiency yielded a small decrease in the herbivore thermal niche along with an increase in system stability (Fig. S5C).

To summarize, these results show that the requirements for nutrients and carbon at temperature extremes have different consequences for community properties. The thermal breadth of herbivore persistence tended to be more strongly restricted by increased herbivore phosphorus demands than by increased herbivore carbon demands (Fig. 3, Fig. S4A). System stability exhibited qualitatively different responses to scenarios: increased herbivore nutrient requirements decreased stability at thermal extremes, whereas increased carbon requirements increased stability at thermal extremes (Fig. 3, Fig. S4B). Increases in both the phosphorus or carbon demands for herbivore growth often decreased interaction strengths, but these decreases occur over different temperature ranges (near the persistence boundary when nutrient demands are increased, and across much of the thermal breadth under increased carbon demands, Fig. 3, Fig. S4C). We next detail the mechanisms by which the temperature-dependence of herbivore nutrient and carbon requirements led to these changes to the herbivore persistence boundary, stability boundary, and interaction strengths.

### Persistence boundary

Increases in herbivore nutrient demands decreases the thermal breadth of the herbivore persistence boundary more strongly than increases in herbivore carbon demands (Fig. 3, Fig. S4A). For example, at a phosphorus supply of *N_tot_* = 0.002 gP m^−3^ and when the herbivore has constant nutrient requirements, the herbivore’s thermal niche spans ∼8.57℃ (see Fig. 4C, where the herbivore thermal niche is demarcated by the vertical dashed lines). Increases in the nutrient or carbon demands of herbivore growth restricts the breadth of the herbivore thermal niche to ∼7.12℃ (Fig. 4F) and ∼8.09℃ (Fig. 4I), respectively.

**Fig. 4.**
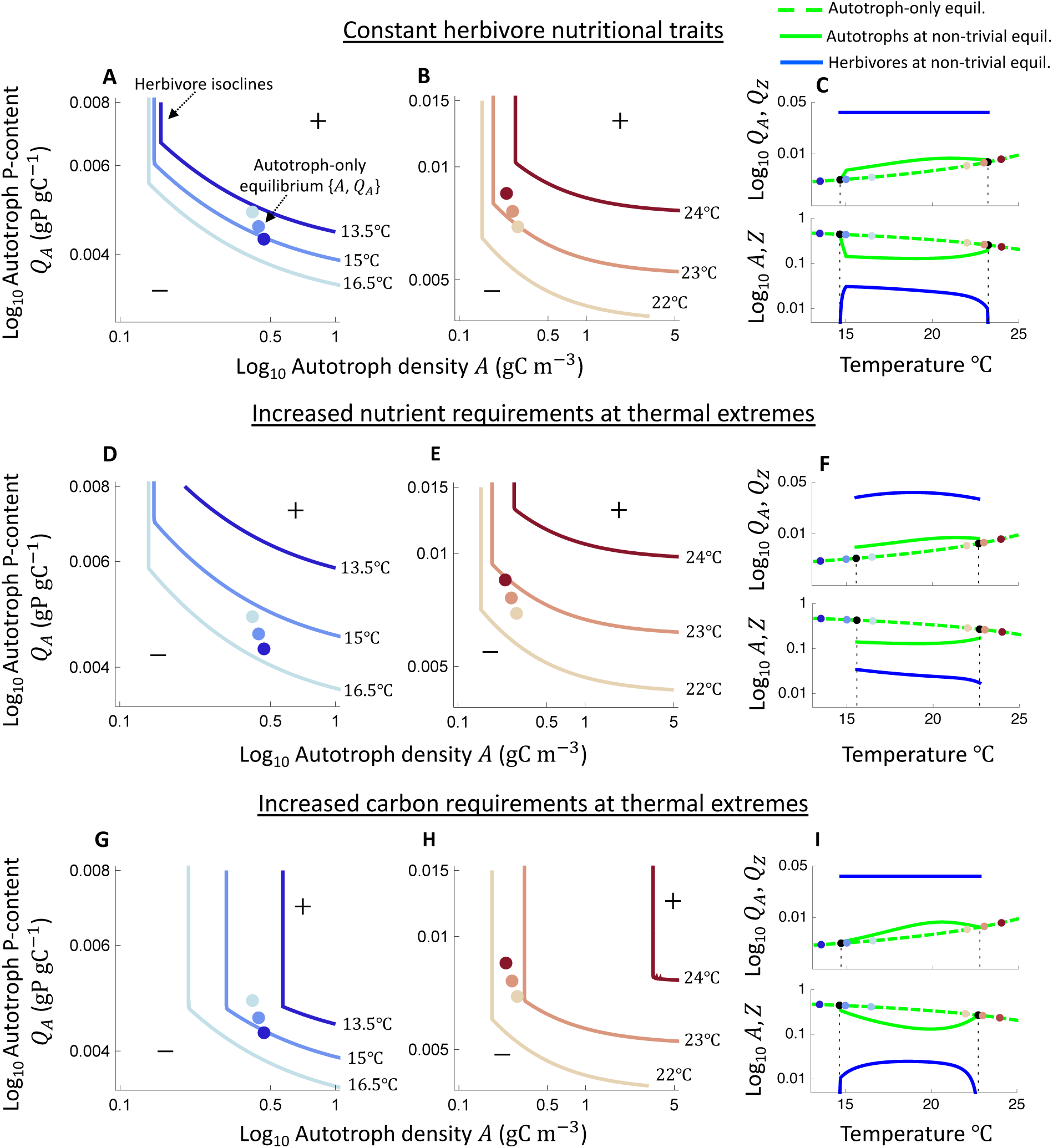
Effects of the temperature-dependence of herbivore nutritional requirements on herbivore persistence. **A-B, D-E**, and **G-H** Effects of temperature on the set of autotroph phosphorus contents and autotroph population densities that allow herbivores to achieve a zero per-capita population growth rate (isoclines). These isoclines are L-shaped because herbivores require a larger quantity of autotroph biomass to maintain zero net growth when autotrophs are of low nutritional quality (autotroph population densities and P-contents above and below these isoclines yield a positive and negative growth rate, respectively). The points in each figure show the autotroph population density and phosphorus content at the autotroph-only equilibrium at each temperature. Importantly, the herbivore population can persist at a given temperature if the autotroph-only equilibrium P-content and density permits a positive per-capita population growth rate (Fig. S3). **C, F,** and **I** Effects of temperature on the phosphorus contents of the autotroph and herbivore at the autotroph-only and non-trivial equilibria (top panel), and on the population densities of the autotroph and herbivore at the autotroph-only and non-trivial equilibria (bottom panels). These panels are included to support interpretation of panels **A-B, D-E,** and **G-H**; the vertical dotted line shows the herbivore persistence boundary for the respective scenario whereas the colored points indicate the autotroph density and P-contents at temperatures considered in **A-B** and **D-E**. This analysis assumes *N_tot_* = 0.002.

We first examine why increased nutrient demands restricts the herbivore’s thermal niche. Consider the impact of temperature on the set of autotroph population densities and phosphorus contents that allow herbivores to achieve a zero net per capita population growth rate under constant (Fig. 4A-B) and increased (Fig. 4D-E) herbivore nutritional requirements. Notice that, at relatively high autotroph population densities (e.g., ∼0.2-1 gC m^−3^), the autotroph P-content that herbivores need to attain a zero net growth rate increases more rapidly with warming and cooling when herbivore nutritional (phosphorus) requirements increase towards warm and cool temperatures (contrast Fig. 4D to Fig. 4A and Fig. 4B to Fig. 4E). To persist, the herbivore must have a positive per-capita population growth rate at the autotroph-only equilibrium population density and phosphorus content (i.e., the colored points in Fig. 4A-B and D-E must lie above the herbivore isocline; see Supplementary Materials section “Calculating community properties” and Fig. S3). At the autotroph-only equilibrium, the autotroph population density and phosphorus content are relatively weakly temperature-dependent (Fig. 4C and Fig. 4F). It is thus the relatively rapid increase in the autotroph P-contents required to support a positive herbivore growth rate that accompanies temperature-dependent increases in nutritional requirements towards thermal extremes that drives the restriction of herbivore persistence boundaries in this scenario.

Increases in the carbon requirements for herbivore growth toward temperature extremes, by contrast, only decreased herbivore persistence slightly relative to the control scenario in which herbivores have constant nutrient requirements. Here, increased carbon requirements decreased the maximum temperature supporting persistence but did not influence the minimum temperature at which herbivores may persist (contrast Fig. 4C with Fig. 4I). Notice that, when autotrophs are relatively P-rich (e.g., >∼0.005 *gP gC*^−1^), increased carbon requirements towards thermal extremes leads to relatively large increases in the autotroph population density that herbivores need to achieve a net zero per-capita population growth rate as temperatures increase (Fig. 4H) or decrease (Fig. 4G). At cold temperatures, autotrophs are at a sufficiently high population density at their equilibrium in the absence of herbivores that this increase in carbon requirements does not influence persistence (contrast Fig. 4G with Fig. 4A). This increase in carbon requirements at warm temperatures did restrict herbivore persistence, however, because autotrophs attain relatively low population densities at the autotroph-only equilibrium (contrast Fig. 4H with Fig. 4B).

Finally, the effects of increasing the herbivore’s relative requirements for phosphorus or carbon at temperature extremes described above (Fig. 3-4) arise because those increased phosphorus or carbon demands reflect impaired phosphorus or carbon growth rates, respectively (Fig. S5). This is evidenced by the analysis in the Supplementary Materials section “Effects of the herbivore *TER_P:C_* on community properties,” which shows that increases in the herbivore’s carbon requirements that emerges from enhanced phosphorus growth rates (i.e., enhanced phosphorus conversion efficiency) expands the herbivore persistence boundary (Fig. S5D) by decreasing the autotroph P-contents that herbivores need to achieve a positive per-capita population growth rate (Fig. S6C-D). Increases in the herbivore’s phosphorus demands that reflect improved carbon growth rates (increased carbon conversion efficiency) has little influence on herbivore persistence (Fig. S5C) because this only influences the autotroph population density that herbivores need to persist, and herbivore persistence is more strongly influenced by autotroph P-contents (Fig. S6A-B).

### Stability boundary

Relative to the scenario with constant herbivore nutrient requirements, increased nutrient requirements generally decreased the phosphorus enrichment needed to destabilize the non-trivial equilibrium whereas increased carbon requirements typically increased it (Fig. 3, Fig. 5, Fig. S4B). These contrasting outcomes reflect different mechanisms. For the increased nutrient requirements at thermal extremes scenario, it is the decrease in herbivore phosphorus contents (*Q*_Z_) and corresponding increase in phosphorus availability for autotrophs that led to decreased stability. Increases in herbivore carbon requirements, however, directly leads to increased stability by decreasing the flux of biomass from autotrophs to herbivores.

**Fig. 5.**
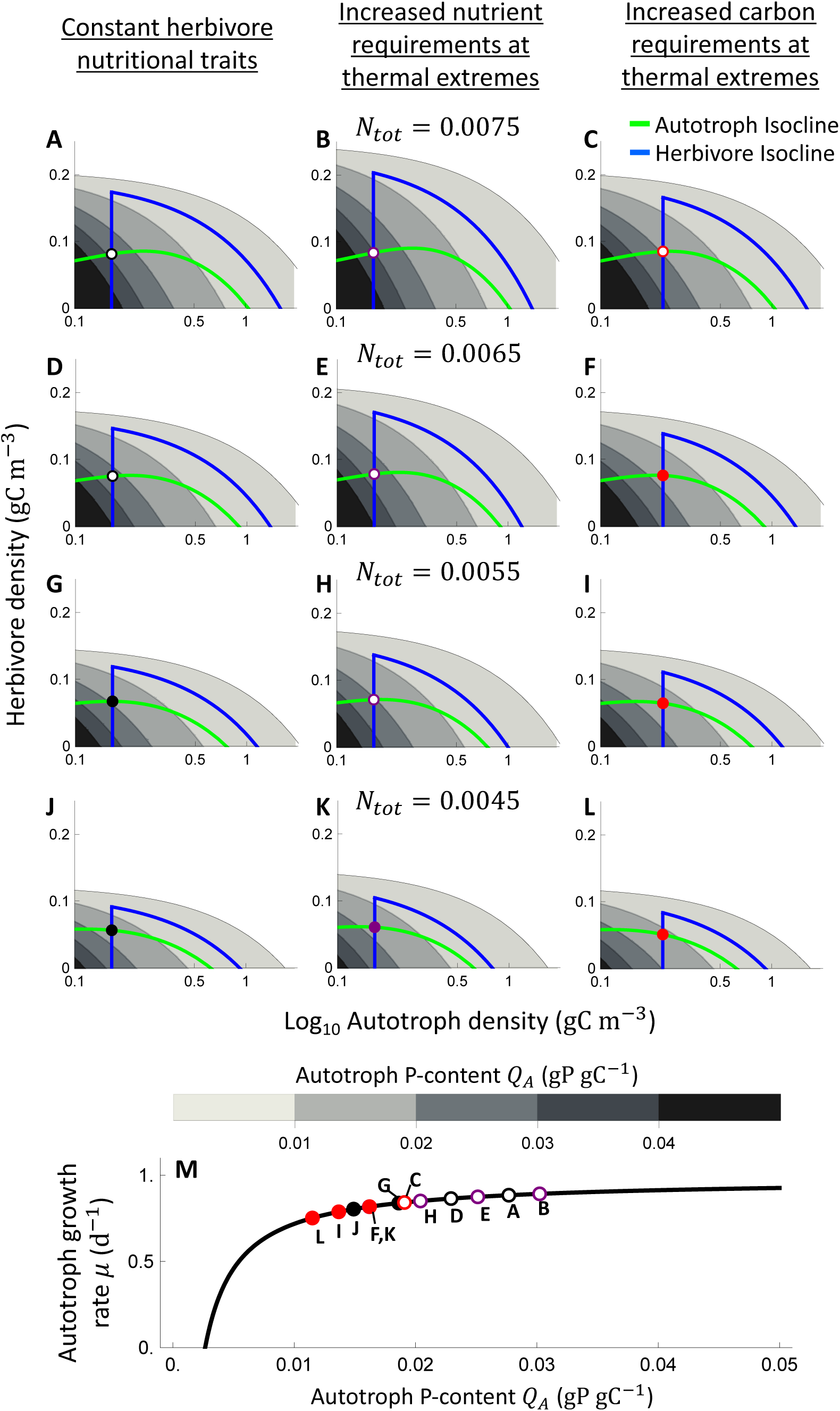
Increases in herbivore nutritional requirements at temperature extremes destabilizes population dynamics at warm (and cold) temperatures, whereas increases in herbivore carbon requirements stabilizes population dynamics. **A-L** Shown are representative isoclines across a gradient in phosphorus availability (*N_tot_* = 0.0075, *N_tot_* = 0.0065, *N_tot_* = 0.0055, or *N_tot_* = 0.0045 for panels **A-C**, **D-F**, **G-I**, and **J-L**, respectively) at a constant temperature (*T*=22.5°C) when herbivores have constant nutritional traits (left column), increased nutrient requirements (middle column), or increased carbon requirements (right column). Importantly, at this temperature, the herbivore in the increased nutritional requirements at thermal extremes scenario has reduced nutritional contents – thus, a given unit of herbivore carbon biomass contains a smaller mass of phosphorus. Herbivores and autotrophs have positive per-capita population growth rates below their isoclines. Unstable and stable non-trivial equilibria are indicated by hollow and solid points, respectively. The gray shading indicates the autotroph phosphorus contents across herbivore and autotroph population density, and white regions indicate densities that violate nutrient conservation (Eqn. S3). The herbivore isocline is unimodal because high herbivore and/or autotroph population densities are associated with a large share of total phosphorus that is fixed in herbivore and/or autotroph biomass – thereby reducing autotroph P-contents, herbivore growth rate, and ultimately herbivore per-capita population growth rate. Notice that instability occurs when the equilibrium lies at densities yielding relatively large autotroph phosphorus contents (*Q_A_* > ∼0.02), where there are locally weak effects of autotroph population density on autotroph growth rate (see panel **I**). **I** Effect of autotroph phosphorus content on autotroph specific growth rate (Eqn. 5) at 22.5°C. The points show the equilibrium phosphorus contents and autotroph birth rates at the indicated panels.

To understand these findings, first recall that an equilibrium is resilient to perturbation (e.g., an increase in autotroph density) if the decline in autotroph growth rate with increased autotroph population density (i.e., a stabilizing mechanism that contributes to a reduction of the autotroph population back towards the equilibrium) is strong relative to the decline in autotroph mortality rate with increased autotroph population density that arises from saturation of the herbivore functional response (i.e., a destabilizing mechanism that contributes to growth of the autotroph population away from the equilibrium). Sufficient increases in phosphorus availability *N_tot_* always destabilizes the equilibrium by concentrating phosphorus in autotroph biomass and thus rendering autotroph growth rate relatively insensitive to autotroph population density near the equilibrium (notice in Fig. 5M that autotroph birth rate is a saturating function of phosphorus contents *Q_A_*, such that autotroph self-regulation is weak at equilibria associated with high *Q_A_*; e.g., Fig. 5A). Note that, although autotroph phosphorus contents *Q_A_* also influences herbivore phosphorus growth rate (*g_P_*(*A*, *Q_A_*, *T*), Eqn. 2c), this does not influence interaction strengths or stability because herbivore growth is carbon-limited and thus not sensitive to *Q_A_* at the non-trivial equilibrium (i.e., the autotroph isocline intersects the herbivore isocline in the vertical portion of the herbivore isocline, Fig. 5A-L).

When herbivores have relatively low phosphorus contents (e.g., at *T*=22.5°C under the increased nutritional requirements scenario), a given unit of herbivore carbon biomass is tied to a smaller mass of phosphorus, thereby increasing the amount of total phosphorus in autotroph biomass (contrast gray contours in Fig. 5B, E, H, and K to those in Fig. 5A, D, G, and J). This mechanism reduces the phosphorus enrichment required to weaken the stabilizing influence of negative density dependence sufficiently to destabilize the equilibrium. Note, too, that changes in the herbivore’s phosphorus conversion efficiency do not influence stability (Supplemental Materials section “Effects of herbivore *TER_P:C_* on community properties,” Fig. S5D) because these changes do not affect autotroph density or nutrient contents at the non-trivial equilibrium (Fig. S7B). Thus, it is the decrease in herbivore nutritional contents *Q*_Z_ towards thermal extremes that ultimately makes the system more sensitive to the destabilizing influence of nutrient enrichment.

Increases in herbivore carbon requirements, however, increases stability by decreasing the herbivore’s capacity to control autotroph population density. Notice that the autotroph population density needed to support a positive per-capita population growth rate of herbivores is increased when herbivore carbon requirements are higher (i.e., the vertical portion of the herbivore isocline is shifted to higher autotroph densities in Fig. 5C, F, I, and L relative to Fig. 5A, D, G, and J). As such, at the non-trivial equilibrium, autotrophs are at a relatively high population density and have correspondingly low phosphorus contents, which strengthens autotroph self-regulation and thereby increases the phosphorus enrichment needed to destabilize dynamics. It is therefore the decreased trophic interaction strength that results from increased herbivore carbon requirements that increases stability.

### Interaction strength

The trophic interaction strength, defined as the reduction in autotroph biomass that occurs in the presence of the herbivore (Eqn. S5), exhibits a hump-shaped relationship to temperature in each scenario (Fig. 6A). The interaction strength always approaches 1, which indicates that herbivores do not displace autotrophs from their equilibrium density achieved in the absence of herbivores, at the herbivore persistence boundaries. Increases in the herbivore’s requirements for either phosphorus or carbon can decrease interaction strengths relative to the control case with constant herbivore nutrient demands, but these decreases occur over different temperature ranges: increased phosphorus requirements decrease interaction strengths at thermal extremes near the herbivore persistence boundary (contrast dashed purple and solid black lines in Fig. 6A), whereas increased carbon requirements decreased interaction strengths at most temperatures (contrast dotted red and solid black lines, Fig. 6A).

**Fig. 6.**
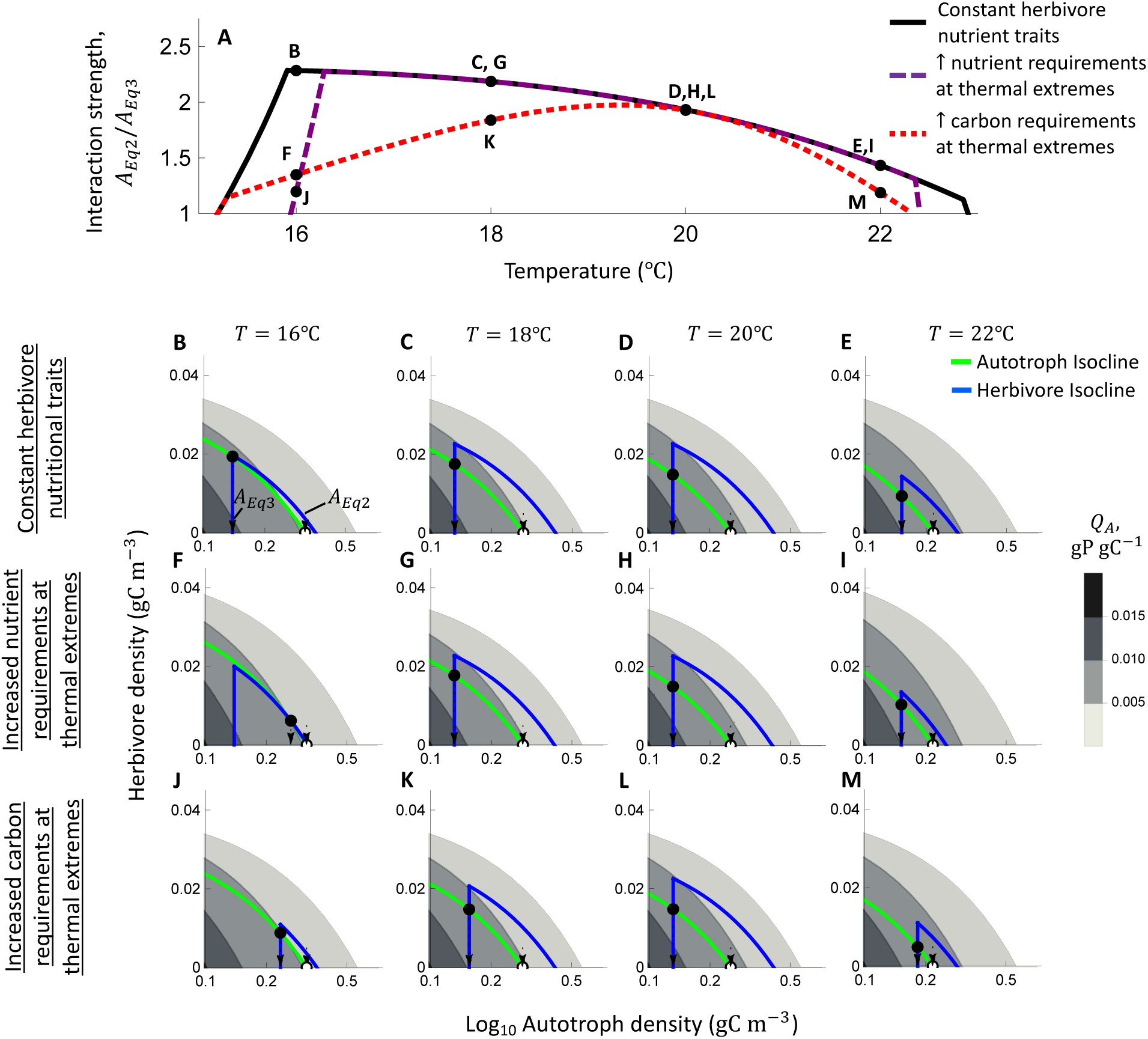
Effects of increases in herbivore nutritional or carbon requirements at temperature extremes on the temperature-dependence of trophic interaction strength. **A** Effect of temperature on trophic interaction strength at a constant phosphorus availability (*N_tot_* = 0.0015 gP m^−3^) when herbivores have constant nutritional requirements (solid black line), increased nutrient requirements at temperature extremes (dashed purple line), and increased carbon requirements at thermal extremes (dotted red line). This figure is a horizontal slice through the contour plots shown in Fig. 3A-C. The points labeled in **A** are considered in more detail in subsequent panels. **B-M** Representative isoclines across a temperature gradient at a constant phosphorus availability when herbivores have constant nutritional traits (**B-E**), increased nutrient requirements (**F-I**), and increased carbon requirements (**J-M**). In each panel, herbivores and autotrophs have positive per-capita population growth rates below their respective isoclines. The gray shading indicates the autotroph phosphorus contents across herbivore and autotroph population density, and white regions indicate densities that violate nutrient conservation (Eqn. S3). The autotroph-only and non-trivial equilibria are indicated by hollow and solid points, respectively. The trophic interaction strength is the ratio of the autotroph density at the autotroph-only (*A_Eq_*_2_) and non-trivial equilibrium (*A_Eq_*_3_) – thus, the trophic interaction strengths can be visually assessed by examining the horizontal displacement of the non-trivial equilibrium (indicated here by arrows in **B-M**).

The mechanisms underlying these patterns can be understood by examining how temperature influences herbivore and autotroph zero net growth isoclines (Fig. 6B-M). The interaction strength reflects the horizontal displacement of the non-trivial equilibrium (*A_Eq_*_3_) from the autotroph-only equilibrium (*A_Eq_*_2_) (Eqn. S5, Fig. 6B). Notice that interaction strengths are unchanged under the increased nutrient demands scenario at intermediate temperatures (e.g., 18°C, 20°C, 22°C) because the equilibrium occurs at autotroph densities at which herbivore growth is carbon-limited (i.e., at the vertical portion of the herbivore isocline in Fig. 6C-E and Fig. 6G-I) and is thus not impacted by differences in the herbivore’s nutrient requirements for growth. It is only near the herbivore persistence boundary (e.g., 16 °C) where the non-trivial equilibrium occurs at densities at which the growth of the herbivore population is limited by autotroph nutrient contents (i.e., the declining portion of the herbivore isocline, see Fig. 6F) that increased nutrient requirements leads to decreased interaction strength (contrast Fig. 6B to 6F).

In contrast, interaction strengths are decreased across much of the temperature gradient when herbivores have increased carbon requirements scenario because, in this scenario, the autotroph population density needed to support a positive herbivore per-capita population growth rate is relatively high (i.e., the vertical portion of the herbivore isocline is at higher autotroph densities in Fig. 5J, K, and M than in Fig. 5B, C, and E). This ultimately leads to a relatively large equilibrium density of autotrophs when present with herbivores, and therefore weak trophic interaction strengths relative to the scenario in which herbivore nutritional requirements are constant (contrast Fig. 6B-D with Fig. 6F-H).

## Discussion

We examined how the temperature-dependence of herbivore nutrient traits modulates autotroph-herbivore population dynamics across temperature and nutrient supply gradients. Our analysis showed that these effects depend on whether herbivores require more nutrient- or carbon-rich autotrophs at thermal extremes. The herbivore’s thermal niche was decreased both by increases in its nutrient and carbon requirements, but this decrease was larger under increased nutrient requirements (Fig. 3-4). Increases in herbivore carbon demands tended to stabilize dynamics under nutrient enrichment, whereas increased nutrient requirements (when accompanied with decreases in herbivore nutrient contents) destabilized dynamics (Fig. 3, 5). Finally, the herbivore’s capacity to control autotroph population density was generally more strongly decreased by increases in the herbivore’s requirements for carbon than for phosphorus (Fig. 3, 6).

Our study advances on theoretical models describing interactive effects of temperature and resource (nutrient) supply on consumer-resource dynamics (Uszko et al. 2017, Sentis et al. 2022, Vinton and Vasseur 2022) by incorporating stoichiometric constraints and thermal variation in herbivore nutrient traits. Consistent with these past studies, we found that the herbivore (consumer) population persists over some temperature range at which their growth rate exceeds losses to mortality (see e.g., Fig. 2B) and that restricted nutrient supply that decreases autotroph (resource) availability led to decreases in the temperature range supporting herbivore persistence (Fig. 3, Uszko et al. 2017, Sentis et al. 2022, Vinton and Vasseur 2022). Similarly, we found that sufficient increases in nutrient supply always led to unstable dynamics, but that decreases in herbivore productivity at temperature extremes can increase the nutrient enrichment needed to destabilize the system by leading the herbivore to control the autotroph at higher densities (creating a U-shaped thermal response of the stability boundary, Fig. 3, consistent with results based on a type 2 functional response in Uszko et al. 2017, Sentis et al. 2022).

Increasing herbivore nutrient or carbon demands did quantitatively alter community properties, however, by impairing herbivore performance at high or low autotroph densities, respectively. Increased herbivore nutrient requirements relatively strongly decreased the thermal breadth of the herbivore persistence boundary because, near this boundary, autotrophs are abundant and thus poor in nutritional quality (as the available nutrients are spread over a larger biomass). This effect arises, in part, because we considered an autotroph food source with a wide thermal niche (Fig. S1F, Fig. 4C) relative to that of the herbivore (Fig. 2B). The thermal traits of autotroph species are known to vary across species (Thomas et al. 2017, Kremer et al. 2025), and this variability could alter how the herbivore thermal niche responds to its’ nutrient traits. For example, an autotroph population with a narrow thermal niche leading it to attain low densities with correspondingly high nutrient contents at thermal extremes (Lemoine 2019, Bieg and Vasseur 2024) may broaden the thermal niche of herbivores with high nutrient demands. Increases in the herbivore’s carbon demands, on the other hand, relatively strongly increased stability by weakening the herbivore’s capacity to control autotrophs at low densities where the stabilizing effects of autotroph self-regulation are relatively weak (Fig. 5). These effects observed here when the herbivore feeds with a type 2 functional response could be altered if it instead exhibits a type 3 functional response, as has been observed for *A. tonsa* and other copepods (Thor and Wendt 2010, Wollrab and Diehl 2015), because, in this case, weakening interaction strengths can be destabilizing (Uszko et al. 2015, 2017).

Which of the thermal responses of herbivore nutrient requirements considered here are more prevalent? Increases in carbon requirements at cool and warm temperature was shown by Ruiz et al. (2020) to arise simply from a strong decrease in ingestion rates relative to metabolic demands. This mechanism seems likely to be widely relevant because ingestion rates are typically unimodal functions of temperature (Englund et al. 2011) and respiration rates often increase exponentially over the temperature range supporting high ingestion rates (Ruiz et al. 2020). These patterns underlie the “metabolic meltdown” hypothesis, which proposes that thermal limits are generated by an inability to acquire enough energy to cover demands (Huey and Kingsolver 2019). Still, Laspoumaderes et al. (2022) presented contrasting evidence from several species that suggest that nutrient demands are increased at temperature extremes. Their experiments found that the nutrient:carbon ratio of herbivore biomass decreased when individuals were raised at temperature extremes (Fig. S8), which indicates the capacity to incorporate nutrients into biomass was impaired relative to the ability to incorporate carbon. Paradoxically, this indicates that there is an inverse relationship between the estimated nutrient requirements for biomass production (*TER_P:C_*) and the nutrient contents of that biomass. The prevalence of this inverse relationship between nutrient requirements and contents across herbivore species is unclear in large part because of insufficient data, but also because a mechanistic explanation of this pattern based on the dynamics of nutrient and carbon uptake and allocation (e.g., Sterner 1997, Sousa et al. 2010, Anderson et al. 2017, Prater et al. 2024) has not yet been developed.

We considered autotroph-herbivore interactions in static environments, yet natural environments are characterized by temporal variability in temperature and nutrient supply. To understand how stoichiometric constraints might affect population dynamics in variable environments, it will be important to consider the timescales at which herbivore nutritional requirements and autotroph nutritional contents respond to environmental change. Herbivores may respond to a nutrient-poor diet by investing resources in enzymes or other digestive traits that aid in extracting nutrients from food (Karasov et al. 2011) or by shifting body composition (Hood and Sterner 2010). These physiological responses can mediate performance in environments with temporal variability in food nutritional quality or temperature (Hood and Sterner 2010, Koussoroplis et al. 2019, Van Baelen et al. 2024). It is well-established that the nutritional quality of autotrophs such as phytoplankton declines with increasing temperature and decreasing nutrient supply (Yvon-Durocher et al. 2015, Tanioka and Matsumoto 2020, Sauterey and Ward 2022), and that the time-scales of these responses have important implications for population dynamics of autotrophs (Caperon 1968, Droop 1974, Bieg and Vasseur 2024, Anderson et al. 2025) and herbivores (Diehl et al. 2022). Future work may consider how flexibility in consumer nutritional requirements interacts with plasticity in autotroph stoichiometry to moderate community structure and dynamics in temporally variable environments.

Resource enrichment can arise both from an increase in nutrient availability, as considered in our model, and from an increase in solar energy (light) supply. How might variation in irradiance, and associated changes in autotroph growth and stoichiometry, alter the response of communities to temperature gradients? In contrast to nutrient-limited scenarios in which herbivores influence autotroph growth by retaining nutrients, autotroph growth is not influenced by the herbivore’s density or stoichiometry under light-limitation (Klauschies and Gaedke 2020). Autotroph stoichiometry also exhibits fundamentally different responses to light enrichment than to nutrient enrichment – increased irradiance leads to decreases in autotroph nutritional contents (i.e., decreased Nutrient:Carbon ratios, as high irradiance enhances carbon fixation), whereas increased nutrient supply increases autotroph nutrient contents (Urabe and Sterner 1996, Tanioka and Matsumoto 2020). This mechanism leads to qualitative changes in the system’s dynamical response to enrichment: increased irradiance initially destabilizes the system (via the classic paradox of enrichment mechanism (Rosenzweig 1971); also observed here with nutrient enrichment, see Fig. 5), but further increases in irradiance and concomitant reductions in autotroph nutritional quality can stabilize population dynamics and ultimately lead to herbivore extinction (referred to as the “paradox of energy enrichment;” (Loladze et al. 2000, Diehl 2007, Elser et al. 2012)). To our knowledge, effects of temperature on properties of the paradox of energy enrichment (e.g., the irradiance needed for stabilization and for herbivore extinction) have not been investigated. We might expect that the irradiance yielding herbivore extinction exhibits a unimodal relationship to temperature, as temperature extremes impair herbivore productivity (via reduced consumption rates (Uszko et al. 2017), increased metabolic requirements (Rall et al. 2010), and/or reduced nutrient conversion efficiency as considered here).

## Conclusion

Nutrients and temperature are key global change factors that influence ecological communities through their effects on organismal stoichiometry and metabolism. In this study, we evaluated the consequences of temperature-dependent changes in herbivore nutritional traits for population dynamics and persistence across gradients in temperature and nutrient supply. We found that increased herbivore nutrient demands and decreased herbivore nutrient content towards temperature extremes restricted the thermal niche of herbivores and altered the cycling of nutrients between herbivore and autotroph populations, which destabilized population dynamics. In contrast, increases in the *carbon* requirements of herbivore growth at temperature extremes slightly decreased the herbivore’s thermal niche, but relatively strongly decreased interaction strengths and thereby stabilized population dynamics. This work advances the understanding of the role of stoichiometry and energetics in moderating the response of population dynamics and persistence to changes in temperature and nutrient availability.

## Supporting information

All supplemental files.

## Notes

### Competing Interest Statement

The authors have declared no competing interest.

### Summary of Updates

Revised the herbivore growth function, and considered effects of increased herbivore carbon demands at temperature extremes.

### Funding

- Simons Foundation: 877200

