## Supplementary material for "Temperature-dependent herbivore nutritional traits affect population dynamics and persistence": All supplemental files.

**Table of Contents:**

- Fig. S1, pg. 2
- Table S1, pg. 4
- Fig. S2, pg. 6
- Supporting text "Calculating community properties," pg. 8
- Fig. S3, pg. 11
- Fig. S4, pg. 13
- Supporting text "Effects of herbivore $TER_{P:C}$ on community properties," pg. 15
- Fig. S5, pg. 17
- Fig. S6, pg. 19
- Fig. S7, pg. 22
- Fig. S8, pg. 24

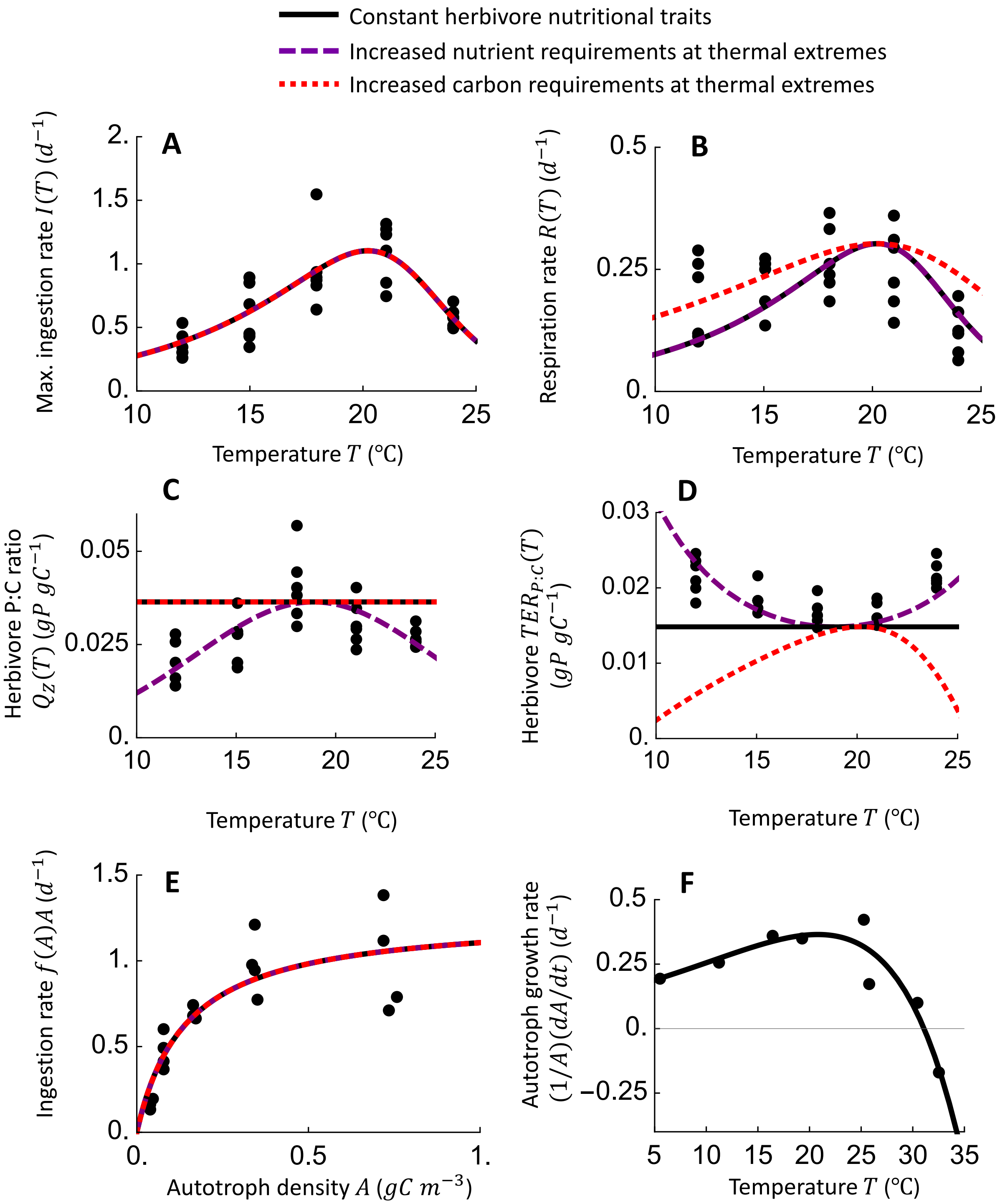


**Fig S1. Fits of model parameters to experimental data. A-B** Fit of Eqn. 3c-d to data for the maximum ingestion rate and respiration rate of *Acartia tonsa* across a temperature gradient from Laspoumaderes et al. (2022). These fits, with the exception of the red dotted line in **B**, were obtained by pooling ingestion and respiration rates and estimating a common set of thermal sensitivities (i.e., assuming $E_{aI} = E_{aR}$, $E_{dI} = E_{dR}$, and $T_{optI} = T_{optR}$) along with the prefactors $I_0$ and $R_0$. The red dotted line in **B** assumes that $E_{aR} = 0.65$ and $E_{dR} = 3$, and $R_0$ was adjusted so that respiration rates are equivalent at $T_{optR}$ across scenarios. Estimates of respiration rate in $gO_2\ gC^{-1}\ day^{-1}$ were converted to $gC\ gC^{-1}\ day^{-1}$ using a respiratory quotient of 0.38 $gC\ gO_2{}^{-1}$ (Andersen 1997). **C** The purple dashed line shows the fit of Eqn. 4a to data for the phosphorus contents of *Acartia tonsa* across temperatures from (Laspoumaderes et al. 2022). The solid black and dotted red lines show the phosphorus contents used in the indicated scenarios (i.e., $Q_z(T_{opt,Q})$). **D** Effects of temperature on phosphorus requirements $TER_{P:C}(T)$ (Eqn. 5) for the three scenarios. The dashed purple line is the fits of Eqn. 5 to data for the phosphorus requirements of *Acartia tonsa* across temperatures from Laspoumaderes et al. (2022). For this fit, the only free parameter was $S_\varepsilon$, which controls the thermal sensitivity of phosphorus conversion efficiency (Eqn. 4b). **E** Fit of Eqn. 3a to data for the ingestion rate of *Acartia tonsa* feeding on *Rhodomonas spp.* at various densities from Thor and Wendt (2010). **G** Fit of autotroph per-capita population growth rate, $(1/A)(dA/dt)$, to data for *Rhodomonas salina* in nutrient-replete conditions and in the absence of grazing mortality from Ferreira et al. (2022). In this case, $Z = 0$ and $Q_A \gg Q_{min}$ (so that $\mu(Q_A, T) \approx \mu_\infty(T)$), and thus $(1/A)(dA/dt) = \mu_\infty(T) - m_A(T)$. This fit therefore yields estimates of parameters for $\mu_\infty(T)$ (Eqn. 6b) and $m_A(T)$ (Eqn. 7a).

**Table S1.** Parameter values and sources. For parameters that vary with temperature, we provide the set of parameters controlling the temperature-dependent function (either a Sharp-Schoolfield, Boltzmann-Arhenius, or gaussian function; see the equation provided in the Model Development section of the main text). Parameter values are provided for each of the three scenarios. Scenario 1 corresponds to "Constant herbivore nutrient requirements," scenario 2 is "Increased nutrient requirements at thermal extremes, and scenario 3 is "Increased carbon requirements at thermal extremes." For values labeled "--", we used the values from Scenario 1.

| **Param.** | **Description** | **Eqn.** | **Units** | **Value** *Scenario 1* | **Value** *Scenario 2* | **Value** *Scenario 3* | **Reference** |
|---|---|---|---|---|---|---|---|
| | | | | *Autotroph* | | | |
| $\mu_\infty$ | Maximum birth rate | Eqn. 6b | $\mathrm{d}^{-1}$ | $\mu_0 = 1.18;$ $E_\mu = 0.56;$ | -- | -- | (Ferreira et al. 2022) |
| $Q_{min}$ | Minimum nutrient requirements | | $\mathrm{gP\ gC}^{-1}$ | 0.00266 | -- | -- | (Meunier et al. 2012) |
| $m_A$ | Mortality rate | Eqn. 7a | $\mathrm{d}^{-1}$ | $m_{A0} = 0.85;$ $E_{mA} = 0.98$ | -- | -- | (Ferreira et al. 2022) |
| | | | | *Herbivore* | | | |
| $a$ | Clearance rate | Eqn. 3b | $\mathrm{m}^3\ \mathrm{d}^{-1}\mathrm{gC}^{-1}$ | $a_0 = 30;$ $T_{opta} = 20.2;$ | -- | -- | (Thor and Wendt 2010, |

| | | | | | | | |
|---|---|---|---|---|---|---|---|
| | | | | $E_{aa} = 1.23$; $E_{da} = 4.81$ | | | Laspoumaderes et al. 2022) |
| $I$ | Max. ingestion rate | Eqn. 3c | $\mathrm{d}^{-1}$ | $I_0 = 3.25$; $T_{optI} = 20.2$; $E_{aI} = 1.16$; $E_{dI} = 4.81$ | -- | -- | (Laspoumaderes et al. 2022) |
| $R$ | Respiration rate | Eqn. 3d | $\mathrm{d}^{-1}$ | $R_0 = 0.83$; $T_{optR} = 20.2$; $E_{aR} = 1.16$; $E_{dR} = 4.81$ | -- | $R_0 = 0.58$; $T_{optR} = 20.2$; $E_{aR} = 0.65$; $E_{dR} = 3$ | (Laspoumaderes et al. 2022) |
| $\varepsilon_c$ | Carbon conversion efficiency | | $\mathrm{gC\ gC^{-1}}$ | 0.6 | -- | -- | (Thor and Wendt 2010) |
| $\varepsilon_p$ | Phosphorus conversion efficiency | Eqn. 4b | $\mathrm{gP\ gC^{-1}}$ | $\varepsilon_{p,0} = 0.8$; $T_{opt,\varepsilon} = 18.9$; $S_\varepsilon = 10^4$ | $\varepsilon_{p,0} = 0.8$; $T_{opt,\varepsilon} = 18.9$; $S_\varepsilon = 4.61$ | -- | (Frost et al. 2006, Laspoumaderes et al. 2022) |
| $Q_Z$ | Phosphorus contents | Eqn. 4a | $\mathrm{gP\ gC^{-1}}$ | $Q_{Z0} = 0.036$; $T_{opt,Q} = 18.9$; $S_Q = 10^4$ | $Q_{Z0} = 0.036$; $T_{opt,Q} = 18.9$; $S_Q = 5.97$ | -- | (Laspoumaderes et al. 2022) |
| $m_Z$ | Mortality rate | Eqn. 7b | $\mathrm{d}^{-1}$ | $m_{Z0} = 0.029$; $E_{mZ} = 0.54$ | -- | -- | (Hirst and Kiørboe 2002, Elliott and Tang 2011) |

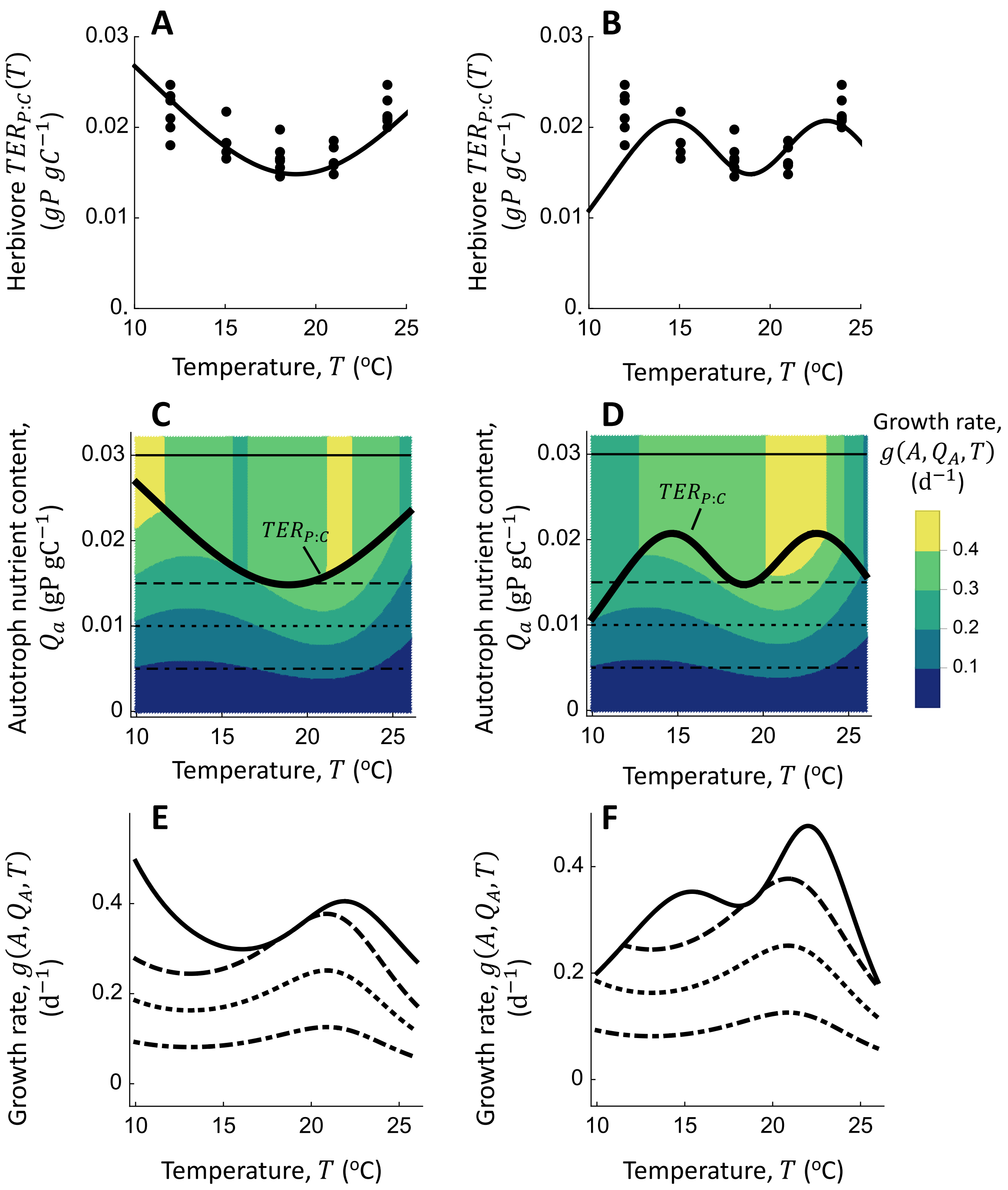


**Fig. S2 Effects of a U-shaped temperature dependence of the carbon conversion efficiency, $\varepsilon_c(T)$, on the temperature-dependence of *TER$_{P:C}$*.** The left column assumes that $\varepsilon_c(T) = \varepsilon_{c0}e^{(T-T_{opt,\varepsilon c})^2/(2S_{\varepsilon c}{}^2)}$, whereas the right column assumes that $\varepsilon_c(T) = \Big(1 -$

$\varepsilon_{cmin} e^{(T-T_{opt,\varepsilon c})^2/(2S_{\varepsilon c}{}^2)}\Big)$. **A-B** These panels show fits of Eqn. 5 to data of the $TER_{P:C}$ of *A. tonsa* from Laspoumaderes et al. (2022) when $\varepsilon_c(T)$ is U-shaped and $\varepsilon_p$ is constant (set at $\varepsilon_p = 0.8$). This analysis estimates the value of the thermal breadth of $\varepsilon_c(T)$, $S_{\varepsilon c}$, with other parameters for $\varepsilon_c(T)$ fixed at $\varepsilon_{c0} = 0.6$, $\varepsilon_{cmin} = 0.4$, $T_{opt,\varepsilon c} = 18.9$. All other parameters underlying Eqn. 5 match those for Scenario 2 in Table S1. **C-D** Effects of autotroph nutrient contents and temperature on growth rate under the two different assumptions for $\varepsilon_c(T)$. The horizontal lines indicate the values of autotroph nutritional quality $Q_A$ considered in panels **E** and **F**. **E-F** Effects of temperature on herbivore growth rate for different values of $Q_A$. These plots assume that autotroph density is not limiting ($A \to \infty$).

*Calculating community properties*

To calculate the community properties considered in this study – the herbivore persistence boundary, the stability boundary, and the trophic interaction strength – we must first define the equilibrium states of the system. Equilibria are defined as the set of autotroph and herbivore population densities (i.e., $\{A_{Eq}, Z_{Eq}\}$) that satisfy:

$$\frac{dZ}{dt} = \frac{dA}{dt} = 0. \quad \text{(S1)}$$

There are three equilibria:

$$Eq_1: trivial, A_{Eq1} = Z_{Eq1} = 0\ , \quad \text{(S2a)}$$

$$Eq_2: autotroph\ only, A_{Eq2} > 0, Z_{Eq2} = 0, \quad \text{(S2b)}$$

$$Eq_3: nontrivial, A_{Eq3} > 0, Z_{Eq3} > 0. \quad \text{(S2c)}$$

To ensure that the mass of phosphorus is conserved, we only consider equilibria with autotroph and herbivore population densities that satisfy (Andersen et al. 2004):

$$Q_{min}A + Q_Z(T)Z \leq N_{tot}. \quad \text{(S3)}$$

To understand this constraint, consider a simple case in which $Z = 0$ and autotroph phosphorus content is fixed at its' minimum possible value $Q_{min}$ ($\mathrm{gP\ gC^{-1}}$). In this case, sufficient increases in autotroph population density $A$ ($\mathrm{gC\ m^{-3}}$) leads the total phosphorus embedded in phytoplankton ($Q_{min}A$; $\mathrm{gP\ m^{-3}}$) to exceed the total phosphorus in the system $N_{tot}$ ($\mathrm{gP\ m^{-3}}$) – thus violating the law of mass conservation. By restricting dynamics to be within the boundary of Eqn. S3, we avoid this issue (Andersen et al. 2004).

We are now in position to define 1) the herbivore persistence boundary, 2) the stability boundary, and 3) the trophic interaction strength. First, we identify the herbivore persistence boundary by solving for the phosphorus availability $N_{tot}$ at which the herbivore maintains a net zero per-capita population growth rate at the autotroph-only equilibrium (Eqn. S2b; Fig. S3A-B; (Andersen 1997, Uszko et al. 2017)):

$$\frac{1}{Z}\frac{dZ}{dt} = g\left(A_{Eq2}, Q_A, T\right) - m_Z(T) = 0, \quad \text{(S4)}$$
$$Q_A = \frac{N_{tot}}{A_{Eq2}}.$$

We then assess the impact of temperature on the phosphorus availability $N_{tot}$ satisfying Eqn. (S4), yielding the set of phosphorus requirements and temperatures that separate herbivore persistence from extinction. This approach (Eqn. S4) can be understood conceptually by comparing the autotroph-only and non-trivial equilibria near the persistence boundary: as reductions in phosphorus availability drive herbivores to extinction, the autotroph density at the non-trivial equilibrium approaches the autotroph-only equilibrium (Fig. S3A-B).

Second, we examined effects of temperature on the total phosphorus concentration $N_{tot}$ at which the system is destabilized, and on the population fluctuations of limit cycle attractors. To do so, we calculated the eigenvalues associated with the Jacobian matrix evaluated at the non-trivial equilibrium (Eqn. S2c), and solved for the $N_{tot}$ at which the real part of the largest eigenvalue equals zero (i.e., the $N_{tot}$ separating a stable equilibrium with negative eigenvalues, and an unstable equilibrium with at least one positive eigenvalue; Fig. S3A, (Uszko et al. 2017)). Stability can be understood by inspecting herbivore and autotroph zero-net growth isoclines (Fig. S3C-D). An equilibrium is stable if a small increase in autotroph population density leads to negative autotroph growth rates and thus a return to the equilibrium (e.g., (Uszko et al. 2015)). This occurs when the autotroph isocline has a negative slope at the intersection with the herbivore isocline (so long as the intersection occurs when the herbivore isocline is vertical), as shown in the example in Fig. S3C (but not Fig. S3D). Enrichment also leads to limit cycles with very low autotroph population densities (Fig. S3A) that leads autotrophs, and thus the herbivores that depend on autotrophs, susceptible to extinction (Rosenzweig 1971). We further track the $N_{tot}$ producing minimum autotroph population densities of $10^{-2}$ and $10^{-4}$ across temperatures.

Third, we assessed the strength of the trophic interaction, denoted $IS$, as the ratio of autotroph density in the absence of herbivory (Eqn. S2b) to the autotroph density in the presence of the herbivore (Eqn. S2c) (Gilbert et al. 2014):

$$IS = \frac{A_{Eq2}}{A_{Eq3}}. \qquad \text{(S5)}$$

Biologically, Eqn. S5 quantifies the potential density of the autotroph population that is captured by the herbivore (see Fig. S3A and C-D), and is a useful metric because it is readily measured experimentally. To understand variability in the $IS$ in the phase-plane, consider the horizontal displacement of the non-trivial equilibrium ($A_{Eq3}$) from the autotroph-only equilibrium ($A_{Eq2}$) (indicated by arrows in Fig. S3C).

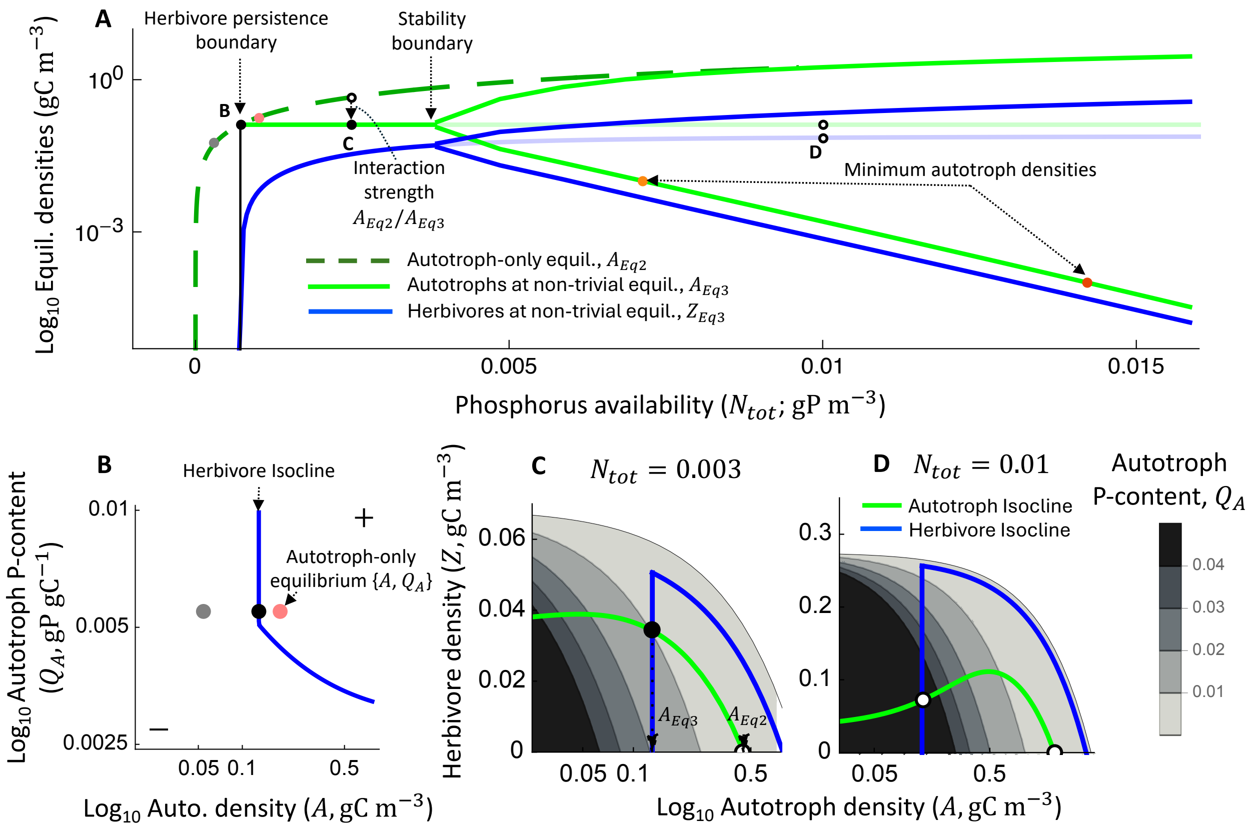


**Fig. S3 Properties of the autotroph-herbivore community considered in this study. A** Phosphorus must be available at a sufficient concentration ($N_{tot}$) to support herbivore persistence, and further enrichment will ultimately destabilize the system and create limit cycles that drive populations to low densities. Shown are the autotroph-only equilibrium population density, and autotroph and herbivore densities at the non-trivial equilibrium. Unstable equilibria are semi-transparent, and in unstable regions we present the minimum and maximum population densities for each population over a limit cycle (opaque lines). The indicated points are explored in more detail in panels **B-D**. **B** Effect of autotroph population density and phosphorus content on the herbivore zero net growth isocline. Herbivores need a larger autotroph population density to persist when autotroph biomass is less nutritious. The gray and pink points indicate autotroph population densities and phosphorus contents at the autotroph-only equilibrium near the herbivore persistence boundary (see **A**). Note that herbivores can persist at an $N_{tot}$ (and temperature) if the autotroph-only equilibrium permits positive growth (as in the pink point). **C-D** Representative isoclines (i.e., sets of autotroph and herbivore densities yielding zero per-capita population growth rates) for a stable (**C**) and an unstable non-trivial equilibrium (**D**). In each panel, herbivores and autotrophs have positive per-capita population growth rates at points below their respective isoclines. The gray shading indicates autotroph phosphorus contents as a function of herbivore and autotroph population density (Eqn. 1c), whereas white regions are densities that violate nutrient conservation (Eqn. S3). Note that herbivores require a greater quantity of autotroph biomass to achieve a positive population growth rate when herbivores are at a high population density (as increasing herbivore density leads to a higher share of total phosphorus locked in herbivore biomass and thus P-poor autotroph biomass – ultimately reducing herbivore assimilation efficiency), creating a unimodal shape of their isocline in phase space. This figure assumes $T = T_{opt,\varepsilon} = 18.9$°C, which is the temperature at which the Constant herbivore nutrient requirements scenario is equivalent to the Increased nutritional requirements at thermal extremes scenario.

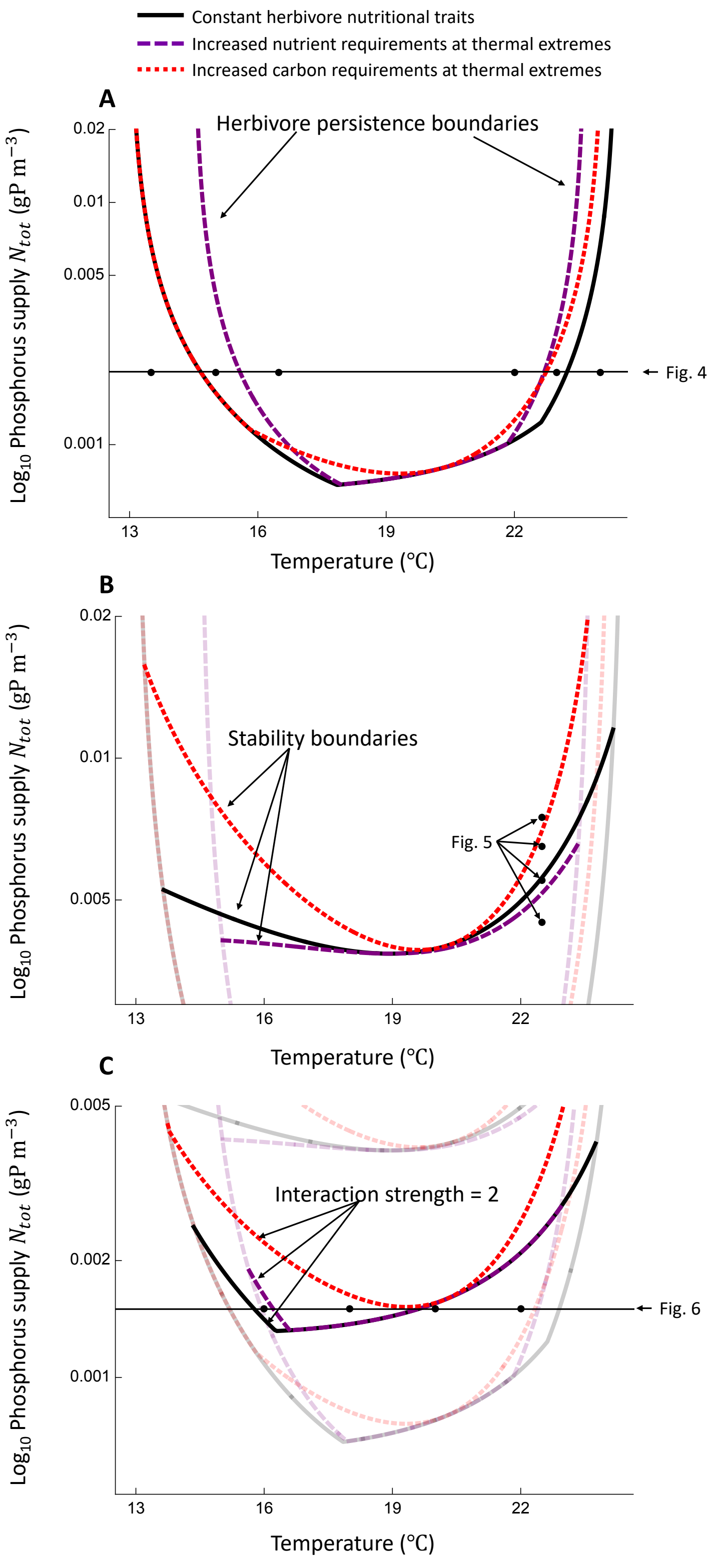


161

162 **Fig. S4 Effects of temperature and phosphorus supply on the herbivore persistence boundary**

**(A), stability boundary (B), and the contour delimiting a trophic interaction strength of 2 (C) for the scenarios in which herbivores have constant nutrient requirements (solid black lines), increased nutrient requirements at temperature extremes (dashed purple lines), and increased carbon requirements at temperature extremes (dotted red lines).** This figure displays the same results as presented in Fig. 3 of the main text, but here the boundaries and contours from the three scenarios are presented together to highlight the differences. The points and thin black lines show the phosphorus supplies and temperatures that are considered in greater detail in figures 4-6 of the main text.

*Effects of the herbivore $TER_{P:C}$ on community properties*

In this section, we evaluate how the herbivore $TER_{P:C}$ influences community properties in the control scenario in which the $TER_{P:C}$ is invariant with respect to temperature (Fig. S5). To do so, we compare the baseline herbivore $TER_{P:C}$ considered in the main text (solid black lines in Fig. S5) to scenarios in which herbivore phosphorus requirements are increased across temperatures due to either an increase in carbon conversion efficiency (dotted cyan lines in Fig. S5) or decrease in phosphorus conversion efficiency (dashed gray lines in Fig. S5), and to scenarios in which the carbon requirements of herbivore growth are increased across the temperature gradient because carbon conversion efficiency is decreased (dashed brown lines in Fig. S5) or because phosphorus conversion efficiency is increased (dotted blue lines in Fig. S5).

These analyses show that the effects of decreasing the herbivore $TER_{P:C}$ on herbivore persistence and stability depend on whether those increases reflect impaired carbon growth rates (dashed brown line in Fig. S5A) or enhanced phosphorus growth rates (dotted blue line in Fig. S5B). When decreased herbivore $TER_{P:C}$ emerged from decreased carbon conversion efficiency, the system generally required larger phosphorus supplies to support herbivore persistence and to destabilize the non-trivial equilibrium (compare brown lines to black lines in Fig. S5C). In contrast, when the decrease in herbivore $TER_{P:C}$ occurs because the herbivore has increased ability to convert ingested phosphorus into biomass, the temperature range supporting herbivore persistence is increased while the stability boundary is unchanged (contrast dotted blue lines to black lines in Fig. S5D).

In turn, the consequences of increasing herbivore $TER_{P:C}$ for herbivore persistence and stability differs depending on whether those increases are due to improved carbon growth rates (dotted cyan line in Fig. S5A) or reduced phosphorus growth rates (dashed gray line in Fig. S5B). Increases in the herbivore carbon conversion efficiency that yields increased $TER_{P:C}$ led to a decrease in the phosphorus supply needed to support herbivore persistence at intermediate temperatures (but largely no changes in the thermal breadth of persistence at higher phosphorus supplies), and a decrease in the phosphorus enrichment needed to destabilize dynamics (contrast dotted cyan lines to solid black lines in Fig. S5C). However, increases in the herbivore $TER_{P:C}$ that arises from impaired phosphorus conversion efficiency led to a restriction of the breadth of the temperature range supporting herbivore persistence but no changes in stability (contrast dashed gray lines with solid black lines in Fig. S5D). We next consider the mechanisms leading to these contrasting results.

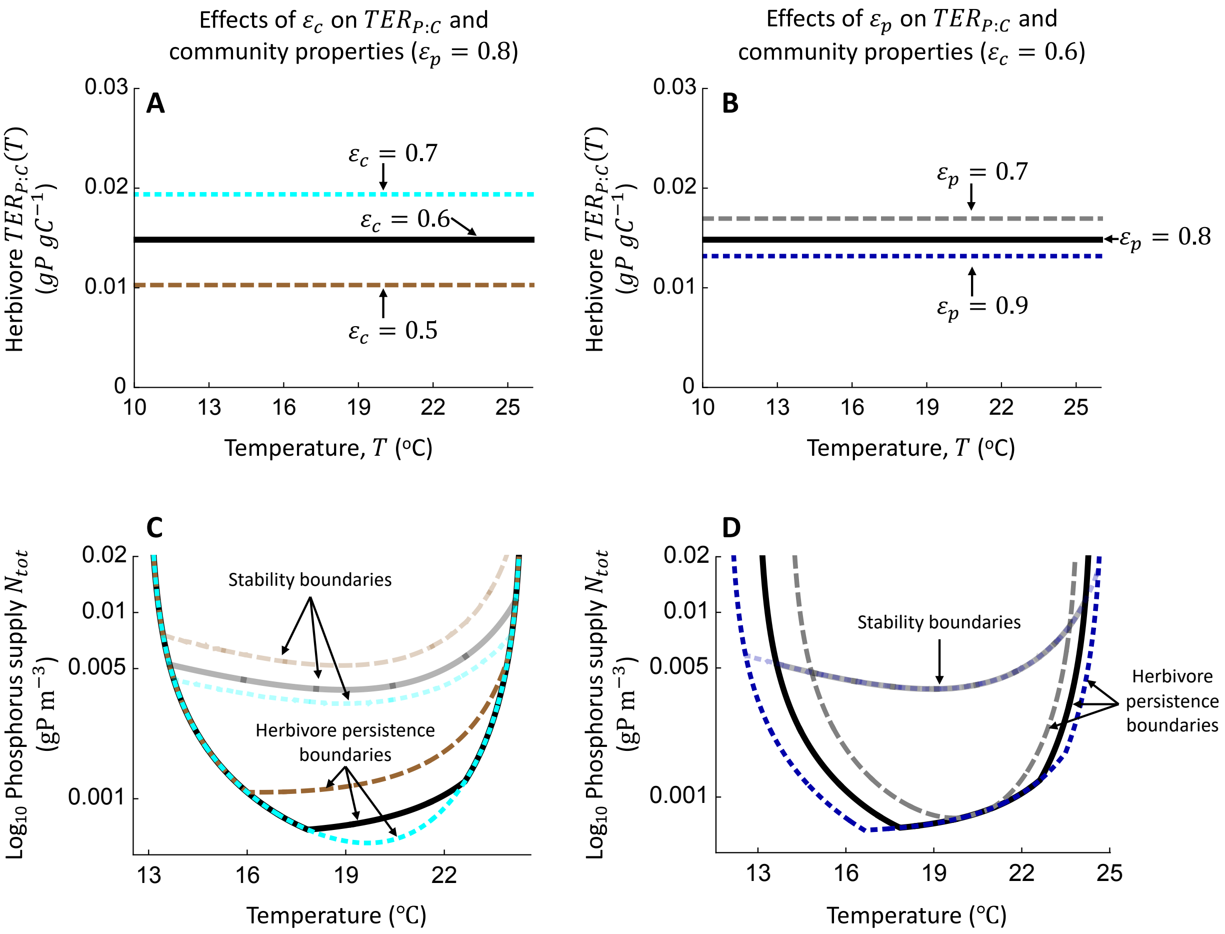


**Fig. S5 Effects of the herbivore *TER*$_{P:C}$ (A-B) on community properties (C-D). A-B** The herbivore *TER*$_{P:C}$ values considered here. The solid black line is equivalent to the "Constant herbivore nutrient requirements" scenario considered in the main text (Fig. 2A-B), whereas the broken colored lines are scenarios involving increased *TER*$_{P:C}$ (due to an increasing carbon conversion efficiency from $\varepsilon_c = 0.6$ to $\varepsilon_c = 0.7$ in **A** or decreasing phosphorus conversion efficiency from $\varepsilon_p = 0.8$ to $\varepsilon_p = 0.7$ in **B**) or decreasing *TER*$_{P:C}$ (due to decreased $\varepsilon_c$ to $\varepsilon_c = 0.5$ in **A** or increasing $\varepsilon_p$ to $\varepsilon_p = 0.9$ in **B**). **C-D** Herbivore persistence boundaries and stability boundaries in temperature—phosphorus supply space for the scenarios indicated in **A-B**. Other parameters are equivalent to those from "Constant herbivore nutrient requirements" in Table S1.

To understand the influence of herbivore $TER_{P:C}$ on herbivore persistence, we examined how the $TER_{P:C}$ affects the set of autotroph population densities and autotroph phosphorus contents supporting a zero-net growth rate of the herbivore population (i.e., the herbivore isoclines, Fig. S6). We focus on the herbivore isoclines at the minimum and maximum temperatures supporting herbivore persistence in the baseline herbivore $TER_{P:C}$ scenario considered in the main text ($T = 14.66$℃ in Fig. S6A, C and $T = 23.23$℃ in Fig. S6B, D, respectively) at a constant nutrient supply ($N_{tot} = 0.002$). Notice that changes in the $TER_{P:C}$ that reflect shifts in carbon conversion efficiency alters the autotroph population density needed to support a positive net growth rate of herbivores when autotrophs are relatively P-rich (i.e., the vertical portion of the herbivore isocline in Fig. S5A-B), whereas changes in the $TER_{P:C}$ that arise from changes in phosphorus conversion efficiency largely influence the autotroph P-contents needed to support net growth of the herbivore when autotrophs populations are abundant (i.e., the approximately horizontal portion of the herbivore iscoline in Fig. S5C-D).

At cold temperatures ($T = 14.66$℃) where the autotrophs are at a high density and have correspondingly low phosphorus contents when they are at equilibrium in the absence of herbivores, changes to the $TER_{P:C}$ due to shifts in phosphorus conversion efficiency do influence persistence (e.g., decreasing $\varepsilon_p$ to $\varepsilon_p = 0.7$ in Fig. S6C leads to herbivore extinction), whereas changes in the $TER_{P:C}$ that reflect shifts in carbon conversion efficiency do not influence persistence (as this does not influence whether the herbivore isocline overlaps the autotroph-only equilibrium, Fig. S6A). At warm temperatures ($T = 23.23$℃), however, decreases in herbivore carbon conversion efficiency do restrict herbivore persistence (contrast dashed brown line with solid black line in Fig. S6B), in part because autotrophs are at relatively low density with correspondingly high nutrient densities at these warmer temperatures.

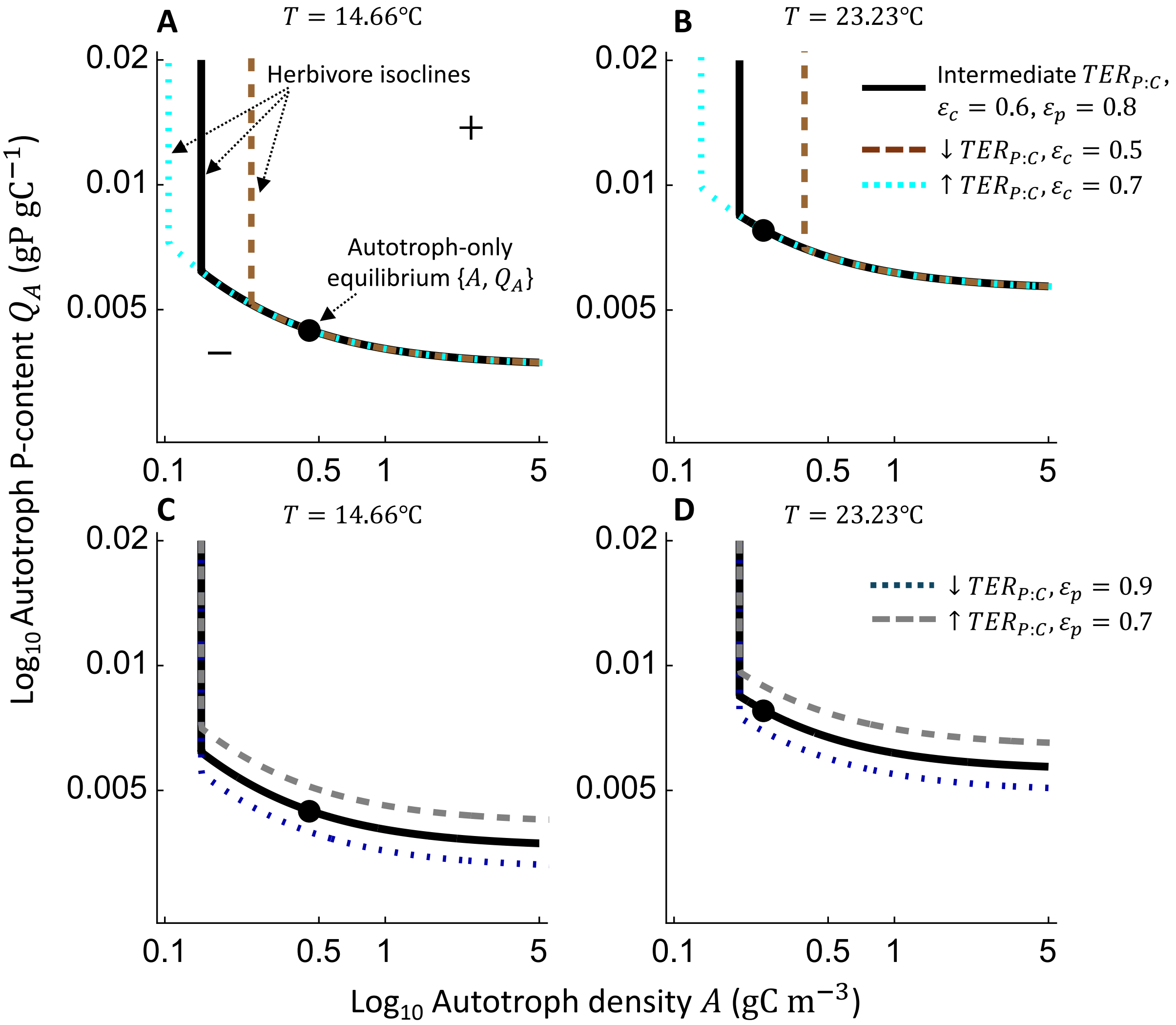


**Fig. S6 The sets of autotroph density and autotroph phosphorus contents that yield a zero-net growth rate of the herbivore population at $T = 14.66$°C (A, C) and $T = 23.23$°C (B, D).**

These temperatures are the minimum and maximum temperatures supporting herbivore persistence, respectively, in the case when $\varepsilon_c = 0.6$ and $\varepsilon_p = 0.8$ and the phosphorus supply is $N_{tot} = 0.002$ (black line). As in Fig. 4 of the main text, the herbivore isoclines are L-shaped because herbivores require a larger quantity of autotroph biomass to maintain zero net growth when autotrophs are of low nutritional quality. Autotroph population densities and P-contents above and below these isoclines yield a positive and negative growth rate, respectively. The points in each figure show the autotroph population density and phosphorus content at the autotroph-only equilibrium at each temperature (note that shifting the $TER_{P:C}$ as we have done here does not influence the autotroph-only equilibrium). Importantly, the herbivore population can persist at a given temperature if the autotroph-only equilibrium P-content and density permits a positive per-capita population growth rate (Fig. S3). The solid black line in all panels is equivalent to the "Constant herbivore nutrient requirements" scenario considered in the main text (Fig. 2A-B), whereas the broken colored lines are the scenarios that increase the $TER_{P:C}$ (reflecting an increased carbon conversion efficiency in **A** or decreased phosphorus conversion efficiency in **B**) or decreasing $TER_{P:C}$ (due to decreased $\varepsilon_c$ in **A** or increased $\varepsilon_p$ in **B**).

The effects of the herbivore $TER_{P:C}$ on the stability of the non-trivial equilibrium can also be understood by examining how changes in the $TER_{P:C}$ influences the herbivore zero-net growth isocline. We do so in Fig. S7 by considering isoclines at the nutrient supply that is needed to destabilize dynamics in the baseline $TER_{P:C}$ scenario at $T = 20°\text{C}$ ($N_{tot} = 0.00394$). Notice that shifts in the $TER_{P:C}$ that reflect changes in carbon conversion efficiency affects system stability by altering the autotroph density needed to support a positive per-capita population growth rate of the herbivore (the vertical portion of the herbivore isocline). For example, decreased $TER_{P:C}$ due to decreased carbon conversion efficiency stabilizes population dynamics in this environment by decreasing the flux of biomass from the autotroph to the herbivore at the non-trivial equilibrium (i.e., by shifting the vertical portion of the herbivore isocline to higher autotroph densities; contrast solid black line to dashed brown line in Fig. S7A). In contrast, shifts in the $TER_{P:C}$ that reflect changes in the phosphorus growth rate influence the autotroph P-contents needed to support herbivore growth (i.e., the declining portion of the herbivore zero-net growth isocline at high herbivore and/or autotroph densities that lead to reductions in autotroph P-contents), and therefore does not influence stability properties of the non-trivial equilibrium.

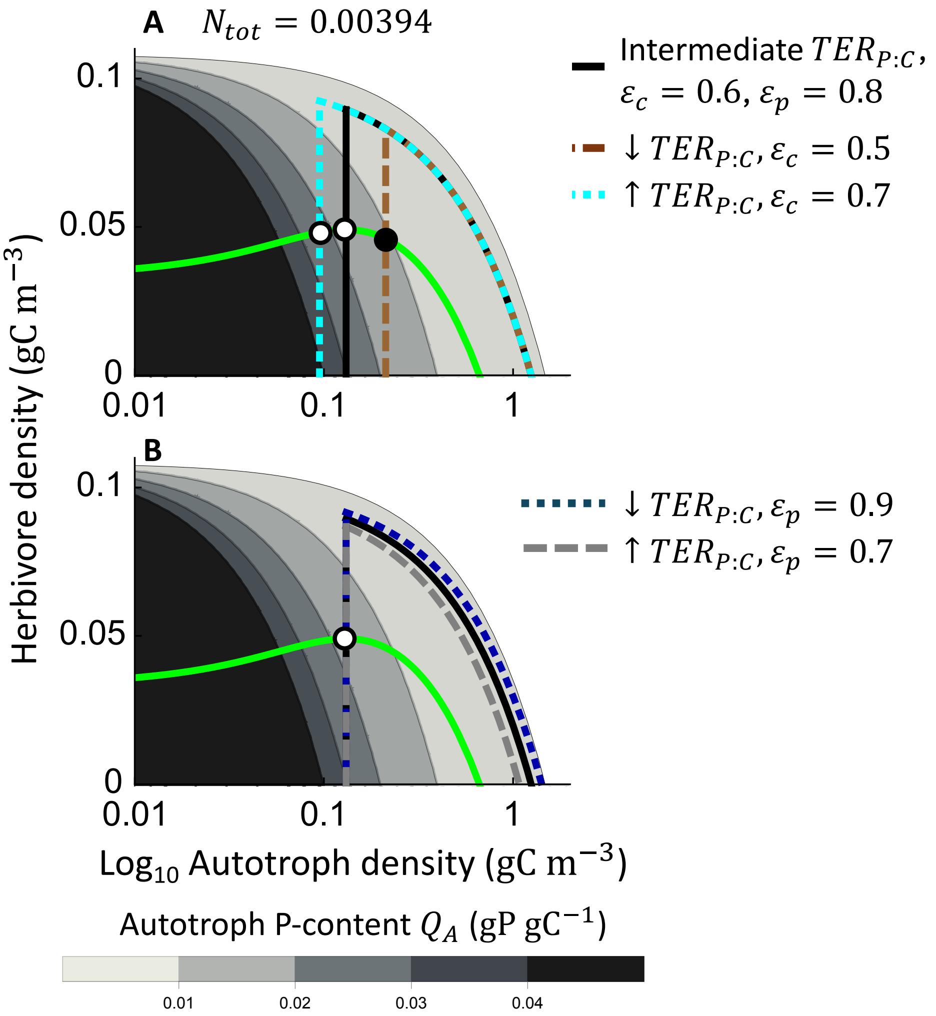


**Fig. S7 Effects of the herbivore *$TER_{P:C}$* on the set of autotroph densities and herbivore densities yielding a zero-net growth rate of the herbivore population.** This analysis assumes that $N_{tot} = 0.00394$, which is the phosphorus supply needed to destabilize the equilibrium when $\varepsilon_c = 0.6$ and $\varepsilon_p = 0.8$ at $T = 20$°C. In both panels, the green line is the autotroph isocline whereas the solid black or broken colored lines are the herbivore isoclines. Both the herbivore and autotroph population have positive per-capita growth rates below their isoclines. Unstable non-trivial equilibria are indicated by hollow points, whereas stable non-trivial equilibria are indicated by solid points. The gray shading indicates the autotroph phosphorus contents across herbivore and autotroph population density, and white regions indicate densities that violate nutrient conservation (Eqn. S3). The herbivore isocline is unimodal because high herbivore and/or autotroph population densities are associated with a large share of total phosphorus that is fixed in herbivore and/or autotroph biomass – thereby reducing autotroph P-contents, herbivore growth rate, and ultimately herbivore per-capita population growth rate. As in Fig. S6, the solid black line in both panels is equivalent to the "Constant herbivore nutrient requirements" scenario considered in the main text (Fig. 2A-B), whereas the broken colored lines include scenarios with increased the $TER_{P:C}$ (reflecting an increased carbon conversion efficiency in **A** or decreased phosphorus conversion efficiency in **B**) or decreased $TER_{P:C}$ (due to decreased $\varepsilon_c$ in **A** or increased $\varepsilon_p$ in **B**).

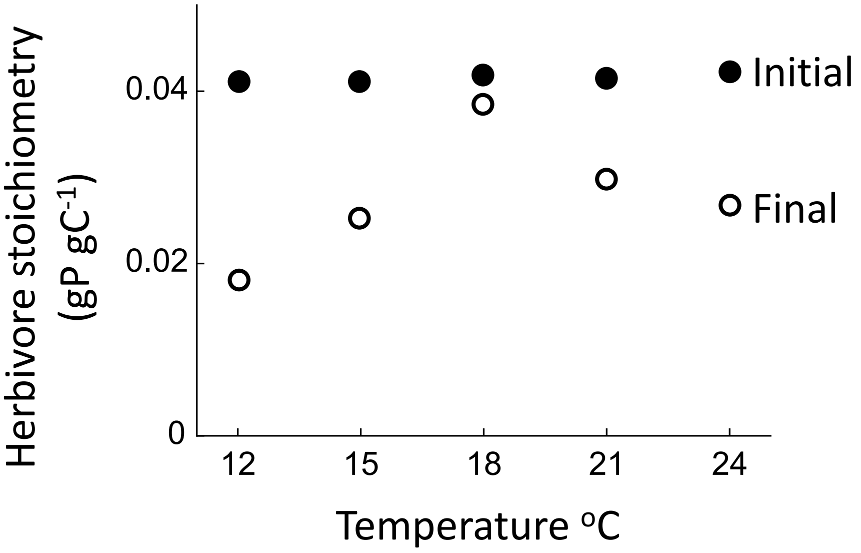


**Fig. S8 Effects of temperature on the body stoichiometry of *A. tonsa* at the beginning (solid points) and end (hollow points) of the experiments conducted by Laspoumaderes et al. (2022).** Data for final body stoichiometry are from Fig. 3B of Laspoumaderes et al. (2022). The initial body stoichiometry is calculated from $TER_{C:P}$ estimates based on initial body stoichiometry in their Appendix 5. $TER_{C:P}$ was estimated by Laspoumaderes et al. (2022) as $TER_{C:P}=(GGE_P/GGE_C)(Q_C/Q_P)$ (their Eqn. 2), where $Q_C/Q_P$ is the carbon:phosphorus ratio of biomass (either initial or final). As such, dividing $TER_{C:P}$ estimates based on initial body stoichiometry in Fig. S5a (panel b) by the ratio $GGE_P/GGE_C$ in Fig. S5b (panel b) (and then converting to units gP gC$^{-1}$) yields the initial body stoichiometry.
